# The Iberian white-oak syngameon as a legacy of introgression in southern Europe

**DOI:** 10.64898/2026.06.09.730892

**Authors:** Carlos Vila-Viçosa, Ricardo Arraiano-Castilho, Francisco M. Vázquez, Rubim Almeida, Cristina García, Albano Beja-Pereira, Andrew L. Hipp, Herlander Azevedo

**Affiliations:** CIBIO, Centro de Investigação em Biodiversidade e Recursos Genéticos, InBIO Laboratório associado, Campus de Vairão; Universidade do Porto, Rua Padre Armando Quintas; 4485-661 Vairão; Portugal; BIOPOLIS Program in Genomics, Biodiversity and Land Planning, CIBIO-. University of Porto Campus de Vairão, 4485-661 Vairão, Portugal; MHNC-UP - Museu de História Natural e da Ciência da Universidade do Porto – Herbário PO, Universidade do Porto. Praça Gomes Teixeira, 4099-002, Porto, Portugal; Department of Biology, Faculty of Sciences, University of Porto, Rua do Campo Alegre, s/n, 4169-007 Porto, Portugal; Department of Ecology and Evolution, University of Lausanne, 1015 Lausanne, Switzerland; Department of Forest Production and Vegetal Biodiversity, Institute of Agricultural Research “Finca La Orden Valdesequera” (CICYTEX), A5 km 372, 06187 Guadajira, Spain; Department of Biological Science. Centre for Evolution, Ecology and Behaviour. Bourne Building, Office 3-29B. Royal Holloway University of London, Egham, Surrey, TW20 EX, United Kingdom; DGAOT, Faculdade de Ciências, Universidade do Porto, Rua Campo Alegre 687, 4169-007 Porto, Portugal; Herbarium and Center for Tree Science, The Morton Arboretum, Lisle IL 60532-1293, USA

**Keywords:** *Quercus* phylogeny; evolution, Mediterranean forests, hybrid swarms, RAD-seq, Roburoid oaks

## Abstract

Oaks (*Quercus* L.) are among the most ecologically important tree genera in the northern hemisphere, with an intricate evolutionary history reflected in a reticulated phylogeny. Oak diversity has been profoundly shaped by introgression and diversification, yet the Iberian Peninsula remains an understudied natural laboratory for understanding these evolutionary processes. We used RAD-seq to characterize 38 taxa (including nothotaxa) and investigate the evolutionary history of the Iberian white oaks, with an emphasis on hybrid swarms. Results led to a readdressing of Iberian white oak species, expanding our current understanding of the phylogeography of the European Section *Quercus*. Furthermore, molecular evidence led to the circumscription of two new subsections, reflecting the contrast between typical temperate and Atlantic distributed species (Group A), and the submediterranean marcescent oaks (Group B). The former unveiled the recovery of *Q. estremadurensis* and a Northwestern Iberian lineage represented by *Q. broteroana* and *Q. orocantabrica* as southwestern representatives of the broad European pedunculate oaks (*Q. robur s.l.*). The latter led to the validation of hybrid swarms, emphasizing the Iberian oak syngameon and the importance of gene flow to oak evolution. Ultimately, our approach advances the understanding of European white oak evolution across different evolutionary scales, establishing the Iberian Peninsula as an important reservoir of oak diversity.

## Introduction

Oaks (*Quercus* L.) are one of the most speciose tree genera in the Northern Hemisphere, where they are keystone elements spanning various forest types and edaphic and climatic niches (Axelrod, 1983; Manos *et al*., 1999; Hipp *et al*., 2020). There are ca. 435 oak species known to date, which display a notoriously complex morphological diversity, stemming from a geographic and ecological diversification of over 56 million years of evolution of wide-range lineages (Denk *et al*., 2017; Carrero *et al*., 2020; Hipp *et al*., 2020). Within Eurasia, the Iberian Peninsula (IP) occupies a unique biogeographic position in Europe and the Mediterranean Basin. Its contrasting bioclimatic gradients (Loidi, 2017) and the presence of a single major biogeographic border to Europe (Pyrenees) turn it into a powerful model for the characterization of refugia and speciation events, associated with past abrupt bioclimatic changes (Williams *et al*., 2011; Veloz *et al*., 2012; Birks & Tinner, 2016).

Section *Quercus*, commonly referred to as the white oaks (Denk *et al*., 2017), includes the Eurasian Roburoid clade, which shows a higher diversification rate throughout the latter half of the Miocene (Hipp *et al*., 2020). The Roburoids present several subclades incorporating temperate and submediterranean distributed taxa, with the latter adapted to the drier Mediterranean climate and with indistinct taxonomic bounderies (Hipp *et al*., 2020). Some of these submediterranean and dry-adapted species have been traditionally assigned to the *Galliferae* subsection or section, depending on the authors (Spach, 1842; Gürke, 1897; Schwarz, 1936b; Schwarz, 1936a). Drought adaptation has evolved convergently in oaks, and together with former Section *Dascia*, *Galliferae* oaks form a notable biogeographic and morphological correspondence within all circummediterranean white oaks, namely in areas where annual and summer precipitations played a significant role in the maintenance of submediterranean marcescent oaks (Schwarz, 1936a; Tschan & Denk, 2012; Gil-Pelegrín et al., 2017; Vila-Viçosa et al., 2020b). However, their relationship with the temperate-dominant species and northern-distributed taxa (*Q. robur* and *Q. petraea*) is yet unclear. Remarkably, the IP harbours almost half of the European oak species and is a well-documented refugium for these taxa (Franco, 1990; Rivas-Martínez & Saénz, 1991; Schwarz, 1993; Vila-Viçosa *et al*., 2020a). This biogeographic positioning of the IP places it as a highly significant yet undervalued model for evolutionary studies.

Sympatric oaks are often not reproductively isolated, and their roles as species syngameons have long been recognized (Van Valen, 1976; Grant, 1981; Hipp, 2015; Hipp *et al*., 2019; Cannon & Petit, 2020; Leroy *et al*., 2020a; Gugger *et al*., 2021; Li *et al*., 2022; Ribicoff *et al*., 2025; Rodríguez-Gómez *et al*., 2025). In syngameons, species tend to remain distinct despite the presence of interspecific gene flow (Cannon & Petit, 2020). Oak taxonomic complexity is reflected at the genome level, with diverse and unique species-level phylogenetic histories, resulting either from introgression or lineage sorting (Eaton *et al*., 2015; McVay *et al*., 2017). In this sense, hybridization complicates the reconstruction of phylogenetic history in an oak syngameon, especially when using traditional organellar DNA markers (e.g., chloroplast or mitochondrial DNA markers) or a combination of a few nuclear loci. Broadly, the lack of resolution in molecular approaches severely hinders the ability to unravel the history of speciation in the reticulated mosaic characteristic of oak genomes (Pham *et al*., 2017; Hipp *et al*., 2020). Thus, there is a need to accurately resolve the phylogenetic backbone of Section *Quercus* as a basis for interpreting its diversification in space and time.

Considering their biogeographical complexity, Iberian white oaks have attracted many taxonomic investigations. *Quercus* species are often difficult to distinguish within broad taxonomic groups due to phenotypic plasticity, genetic variability, and introgression among sympatric species (Schwarz, 1936b). In this regard, studies often lack accurate identification and well-circumscribed species to serve as references. Similarly, the proliferation of names and misidentifications hampers downstream work, from ecological modeling to phylogeny to forest conservation, underscoring the need for state-of-the-art molecular characterization of oak biodiversity (Vila-Viçosa *et al*., 2023). Next-generation sequencing (NGS) technologies and new computational techniques currently provide unprecedented tools for estimating the evolutionary history of living organisms, including oaks (Petit *et al*., 2013; Crowl *et al*., 2020; Leroy *et al*., 2020b; Plomion & Martin, 2020; Backs & Ashley, 2021; Gugger *et al*., 2021). Restriction-site Associated DNA sequencing (RAD-seq) addresses genetic polymorphisms across the genome (Andrews *et al*., 2016), and has been extensively used in studies targeting oak evolutionary history (Hipp *et al*., 2014; Cavender-Bares *et al*., 2015; Eaton *et al*., 2017; Fitz-Gibbon *et al*., 2017; Deng *et al*., 2018; Hipp *et al*., 2020; Lazic *et al*., 2021; Tóth *et al*., 2021; Ulaszewski *et al*., 2021).

The biogeographic significance of the IP in the evolution of Eurasian white oaks remains poorly understood. We hypothesize that the biodiversity of the IP’s white oaks has been consistently undervalued, with the profusion of names and unclear taxonomic assumptions making their phylogenetic resolution urgent. It is also critical to assess the role of the IP as a diversity hotspot within the context of the macro- and microevolutionary processes that shaped the evolutionary history of Eurasian white oaks. In the present study, we addressed the IP’s white oaks, their species and nomenclature uncertainties, by filtering target taxa using extensive literature and fieldwork surveys. We subsequently used RAD-seq to conduct a phylogenomic analysis of key Iberian species, including hybrids, to address species delimitation and phylogenetic relationships among individuals. We analyzed 263 individuals from our sampling set and added 60 individuals from previously published RAD-seq data, representing 38 taxa and nothotaxa (i.e., hybrids) of Iberian and non-Iberian origins. A combination of phylogenetic and population structure methods was used to resolve the backbone of western Mediterranean white oaks within the Eurasian clade of *Roburoid* oaks (Denk et al., 2017; Hipp et al., 2020), thereby developing and extending the role of hybrids in the Iberian syngameon.

## Materials and Methods

### Focal species/groups and study area

We collected 263 white oak samples from the IP (Fig. 1; Method S1). This included 5-10 plants per subpopulation, representing eight species across a west-to-east gradient, which initially targeted *Galliferae* oaks (Gürke, 1897; Schwarz, 1936a; Tschan & Denk, 2012) and reported putative hybrids and related parents within Section *Quercus*. We favored collections in historical type locations, assigned to taxonomic and literature-based nomenclatural endorsements that are considered narrow endemics (e.g., *Q. alpestris*, *Q. orocantabrica* and *Q. pauciradiata*). Our sampling was mostly distributed across the IP’s submediterranean belt, with incursions into Eurosiberian and strictly Mediterranean areas (Vila-Viçosa *et al*., 2020b) (Fig. 1). The Iberian White oaks sampling distribution map was generated in ESRI ArcGIS v10.3 and overlaid with the Iberian life zones (Temperate, Transitional Temperate-Submediterranean, Submediterranean, strictly Mediterranean), based on the SDMs of the deciduous and brevi-deciduous (marcescent) species, which were retrieved by Vila-Viçosa and co-workers (2019; 2020a). A descriptive overview of each taxon and nothotaxon is summarized in Note S1. Several taxa were sampled under narrow taxonomic concepts derived from the historical and regional literature, especially where names were associated with type localities, geographically restricted *populations*, or long-standing Iberian taxonomic usage. Accordingly, *Q. broteroana* and *Q. orocantabrica* were initially treated as distinct sampling labels and are shown separately in the sampling map and analytical outputs. This reflects the original sampling design and field/literature-based identifications, not a final taxonomic conclusion. Subsequently, their evolutionary relationship was evaluated explicitly in both the Results and the Discussion sections.

**Fig. 1.**
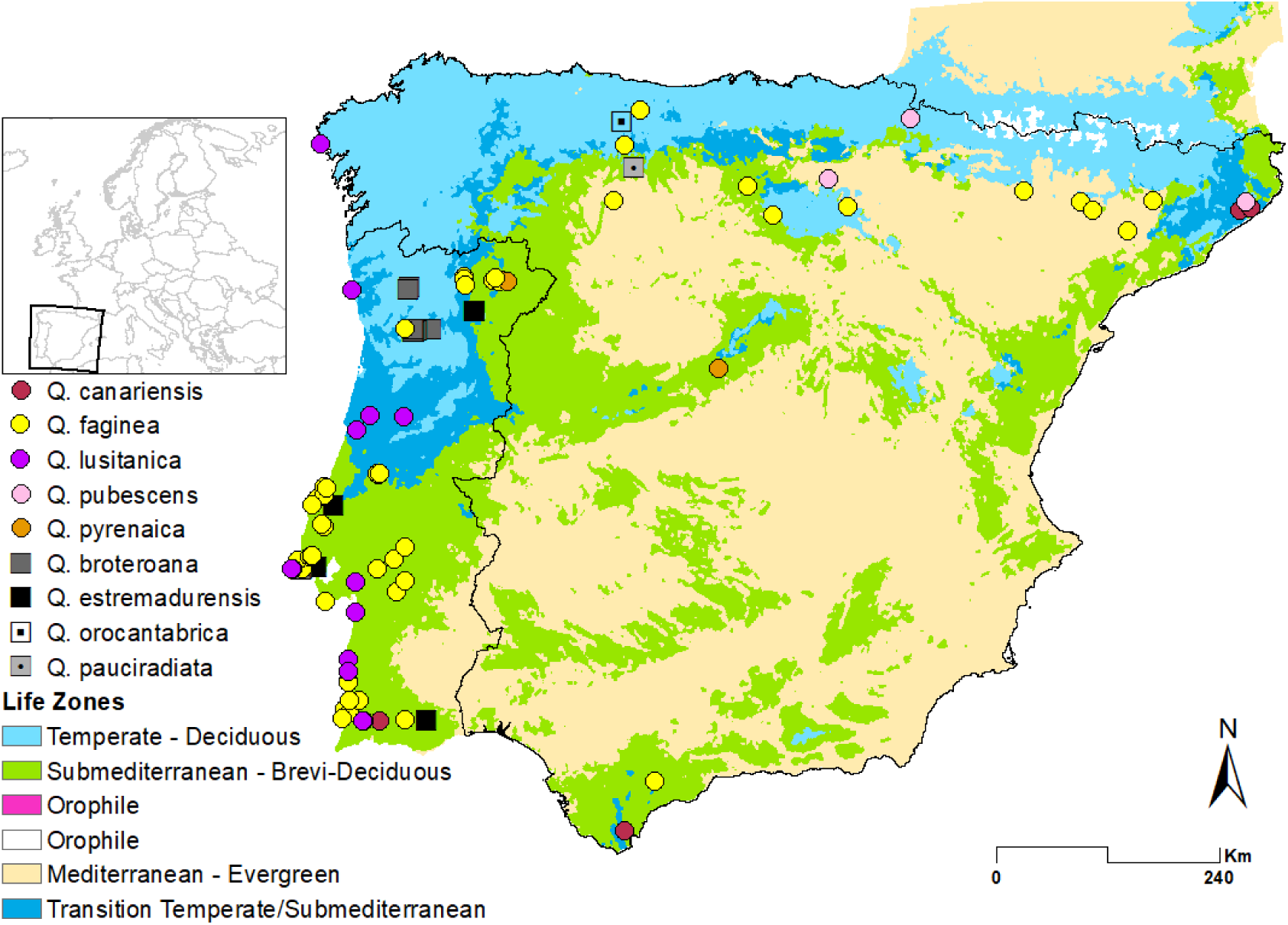
Iberian White oaks sampling distribution. Sampling locations are placed above the main biogeographic lifezones obtained by modelling deciduous (Temperate-Eurosiberian) and brevi-deciduous/marcescent (Submediterranean) oaks, with the overlap of both lifezones considered to be transitional (Eurosiberian-Submediterranean). Squares represent true roburoid oaks and circles represent former Sect. *Dascia* and Sect. *Galliferae* species. Adapted from Vila-Viçosa *et al*. (2019). Taxon labels reflect the original field and literature-based sampling identifications.

### Genomic DNA extraction and sequencing

Genomic DNA (gDNA) was extracted from *ca*. 70 mg of dry or frozen (−80°C) leaf material, using the CTAB-based variation of the NucleoSpin Plant II kit (Macherey-Nagel), according to the manufacturer’s instructions except for an increased (1 h) buffer incubation period. DNA purity, concentration, and integrity were inferred from spectrophotometry (Nanodrop 2000, ThermoFisher Scientific), Qubit dsDNA HS Assay Kit, Picogreen Quant-iT dsDNA Assay kit (Invitrogen, ThermoFisher Scientific), and agarose gel electrophoresis. DNA was normalized to 20 ng/μl in a 50 μl volume for each sample. RAD-seq libraries were prepared using the restriction enzyme *PstI*, barcoded by individual, and sequenced in 100 bp single-end reactions on an Illumina HiSeq 4000 platform at the University of Oregon Genomics and Cell Characterization Core Facility (Eugene, OR, USA) following the RAD-seq protocol of Baird *et al*. (2008).

### RAD-seq loci call and matrix assembly

Raw sequencing reads were processed using *ipyrad* v0.9.31 (Eaton & Overcast, 2020). FASTQ files were demultiplexed and filtered to remove sequences with >5 bases of quality score <20. High-quality reads were clustered at an 85% similarity threshold, with a maximum depth within clusters of 10,000 reads, and mapped back to the *Q. robur* haploid genome assembly PM1N (Plomion *et al*., 2018). All the remaining parameters were set to default, except for the minimum number of samples with data at a given locus, which was set differently for each individual analysis (see below).

### Phylogenetic analysis and species tree inference

To investigate the position of the Iberian clade within the Eurasian white-oaks (Sect. *Quercus*) context, we used a total of 69 samples, including 17 from our field sampling effort and previously published samples representing 38 Eurasian samples and 14 American white oaks (Hipp *et al*., 2020). The RAD-seq loci data matrix was generated using the *ipyrad* assembly previously described and filtered to retain only loci present in a minimum of 52 individuals (*i.e*., 75% of the total number of individuals). The concatenated sequence matrix was assembled from 28,473 filtered RAD-seq loci totaling 5,298,608 base pairs (29.33% missing sites). The maximum likelihood phylogenetic analysis was conducted locally in *RAxML* v.8.2.9 (Stamatakis, 2014) using the GTRCAT implementation of the general time reversible model of nucleotide evolution, with branch support assessed using the RELL bootstrapping method (Minh *et al*., 2013).

To assess the phylogenetic relationships among the Iberian white oaks, we collected 28 representative samples of the currently recognized Iberian clades, which were complemented with previously published data from 22 Eurasian and 3 American white oaks (Hipp *et al*., 2020). The RAD-seq loci data matrix was generated using the *ipyrad* assembly feature described earlier and filtered to include only loci present in at least 40 individuals (*i.e.*, 75% of the total number of individuals). The concatenated sequence matrix was assembled from 30,951 filtered RAD-seq loci totaling 5,702,695 base pairs (29.09% missing sites). The maximum likelihood phylogenetic analysis was conducted in *RAxML* v.8.2.9 as previously described. Species tree inference was based on the SVDQuartets algorithm (Chifman & Kubatko, 2014) using the coalescent-based method tetrad as implemented in *ipyrad*. Briefly, it uses phylogenetic invariants to resolve quartet trees from SNPs for all sets of quartets in a larger tree, and then joins the quartets together into a supertree using the quartets MaxCut algorithm (Snir & Rao, 2012). Here, we subsampled a single SNP from every locus separately (hereafter referred as unlinked SNPs) for every quartet set in the analysis, and repeated this independently over 100 bootstrap replicates. Using this approach, we maximized the amount of unlinked SNP information used in every quartet inference. The average number of unlinked SNPs used to infer each quartet was 27,287 ± 132.

### Genetic assessment using PCA and Neighbor-net analysis

Principal component analysis (PCA), as implemented in the ipyrad tool, was used to evaluate genetic structuring and differentiation among morphologically similar species and their relatives within the Iberian clade. We aligned all samples from each putative subsections inside sect. *Quercus* (Group A and Group B) to the *Q. robur* reference genome (see above). Because several names were deliberately retained as initial sampling labels, we refer below to taxonomic labels rather than final taxonomic entities when describing the analytical matrices. The dataset of Clade Group A was composed of eight initial taxonomic labels (*Q. broteroana, Q. dalechampii, Q. estremadurensis, Q. hartwissiana, Q. orocantabrica, Q. pauciradiata, Q. petraea* and *Q. robur*), retained to test whether field- and literature-based entities corresponded to independently structured genomic lineages. The dataset of Clade Group B was composed of ten initial taxonomic labels (*Q. boissieri, Q. canariensis, Q. faginea, Q. frainetto, Q. lusitanica, Q. macranthera, Q. kotschyana, Q. pubescens, Q. pyrenaica* and *Q. vulcanica*). These datasets incorporated Iberian species as well as their putative hybrids (*Q. ×cerrioides*, *Q. faginea × Q. pyrenaica*, *Q. ×marianica* and *Q.* ×*subpyrenaica*) (Note S1). RAD-seq loci were called individually in each analysis. Only loci observed in at least 75% of the individuals in each of the two datasets were selected. In addition, SNPs present in less than 50% of all the samples in each dataset were discarded, and to reduce the effect of linkage in PCA results, only one SNP was randomly selected per each RAD-seq locus. The SNP subsampling was replicated 25 times to include a different random set of unlinked SNPs, with the total number of selected SNPs per dataset being the following: all species (10,801), Group A (25,021), Group B with hybrids (12,947), Group B without the hybrids (13,535) and all species with hybrids (9,598). To reduce the effects of missing data in all datasets, we tested the imputation methods *sample* and *k-means*, as implemented in *ipyrad*. Both yielded similar results, and for that reason, only the *sample* imputation method results were presented.

Additionally, we inferred two phylogenetic networks based on the maximum-likelihood (GTR+γ) pairwise distance matrix estimated in RAxML. The networks were visualized using the neighbor-net algorithm (Bryant & Moulton, 2004) in *SplitsTree* v4.16.1 (Huson & Bryant, 2006). The first Neighbor-Net included a subset of Asian white oaks (*Q. aliena, Q. dentata*, *Q. yunannensis*, *Q. griffithii*, *Q. fabrii and Q. mongolica*) as outgroup, together with initially identified species and sampling labels (*Q. robur, Q. broteroana, Q. estremadurensis, Q. petraea, Q. pauciradiata, Q. pubescens, Q. macranthera, Q. vulcanica, Q. kotschyana, Q. boissieri, Q. faginea, Q. canariensis, Q. lusitanica,* and *Q. pyrenaica*). The second Neighbor-Net incorporated the studied hybrids (Q. *×cerrioides, Q. ×coutinhoi, Q. ×duriensis, Q. ×fontquerii, Q. ×marianica, Q. ×subpyrenaica* and *Q. faginea × Q. pyrenaica*), among the putative parent species.

### Population structure analysis

To investigate patterns of population structure and hybridization across the main groups identified in the previous analyses, we used the software *STRUCTURE* v2.3 as implemented in the *ipyrad* analysis toolkit. In brief, all structure analyses were run using only SNPs with a minimum allele frequency of 0.05, considering burning periods of 100,000 steps followed by 900,000 additional Markov Chain Monte Carlo iterations, and 10 replicates for each putative cluster (K). The highest deltaK for each Structure run was used to determine the most likely K for each group (Evanno *et al*., 2005).

To assess genetic structure among species within Iberian white oak lineages, we performed STRUCTURE analyses separately for the two major clades previously identified through phylogenomic inference: Group A (temperate roburoid oaks) and Group B (marcescent/submediterranean oaks). For each STRUCTURE run, we used SNP matrices filtered to include only unlinked SNPs with a minimum allele frequency of 0.05 and present in at least 75% of individuals in each dataset. The total number of SNPs used per run was as follows: Group A: 25,021 SNPs; Group B (including hybrids): 12,947 SNPs. This design enabled us to test species boundaries and assess signals of hybridization, while minimizing confounding effects across clades.

## Results

### Phylogenetic analysis of European and Iberian white oaks

In the present work, the challenging phylogeny of European and circummediterranean white oaks was addressed using RAD-seq (Fig. 2), which resolved the species into American, Asian, and European clades, as previously demonstrated (Hipp *et al*., 2020). Notably, the phylogeny revealed two distinct groups within the European white oaks, reflecting the circumscription of two new subsections within the Eurasian species of Section *Quercus,* hereafter designated as Group A and Group B. Group A included roburoid oaks, namely former Sections *Roburoides* and *Robur* (Schwarz, 1936b; Schwarz, 1936a). Additionally, samples and initial taxonomic labels were ascribed to two subgroups, tentatively designated as *Q. robur s.l.* (*Q. robur* L.; *Q. broteroana* (=*Quercus robur* subsp. *broteroana* O.Schwarz); *Q. orocantabrica* Rivas Mart., Penas, T.E.Díaz & Llamas; *Q. estremadurensis* O. Schwarz*; Q. hartwissiana* Steven*; Quercus robur* subsp. *imeretina* (Steven ex Woronow) Menitsky (=*Quercus imeretina* Steven ex Woronow); *Quercus robur* subsp. *pedunculiflora* (K.Koch) Menitsky (=*Q. pedunculiflora* K.Koch) and *Q. robur* L.), and *Q. petraea s.l.* (*Q. dalechampii* Ten. *Q. pauciradiata* Penas, Llamas, Pérez Morales & Acedo; *Q. petraea* (Matt.) Liebl and *Quercus petraea* subsp. *iberica* (Steven ex M.Bieb.) Krassiln (=*Quercus iberica* M.Bieb). Group B comprised all taxa from former sections *Dascia* (*Q. frainetto* Ten.*, Q. kotschyana* O.Schwarz*, Q. pubescens* Willd.; *Q. pyrenaica* Willd. and *Q. vulcanica* Boiss. ex Kotschy) and *Galliferae* (*Q. boissieri* Reut. *Q. canariensis* Willd.*, Q. faginea* Lam. and *Q. lusitanica* Lam.) (Schwarz, 1936a). Here, also, there was a separation into two sub-groups. The first contained an Iberian set of species, also distributed across the Western Mediterranean Basin, including North Africa. The second could be further divided into three subclades, comprising: 1) broadly distributed *Q. pubescens*; 2) species from the Caucasus and Anatolia; 3) species from the Near East.

**Fig. 2.**
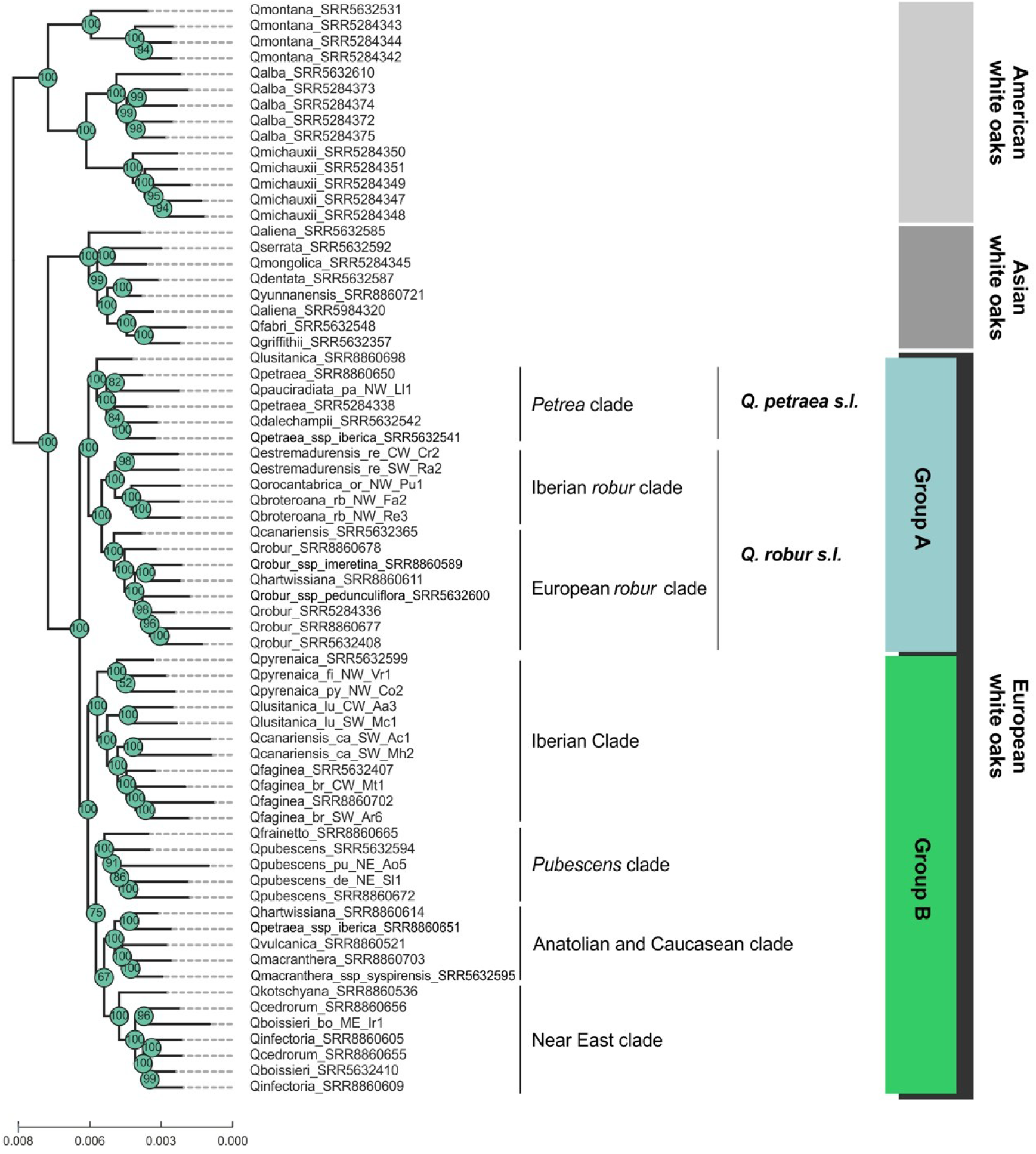
Phylogeny of the Eurasian white-oaks (*Sect. Quercus*), including subsets of American and Asian species used as outgroups.

We focused further on Iberian white Oaks via SVDQuartets analysis (TETRAD test) (Fig. S1). Using American and Asian oaks as outgroups, we observed a segregation of three species: the two existing Anatolian and Caucasian clade members (*Q. macranthera* and *Q. vulcanica*), plus *Q. frainetto*, previously positioned in the *pubescens* clade member (though it presented weaker support in the tetrad tree in Fig. S1B). Samples were further resolved into Groups A and B as before, separating series *Dascia* and *Galliferae* imperfectly. In Group A, samples initially assigned to *Q. broteroana* and *Q. orocantabrica* formed part of the same Northwestern Iberian *Q. robur s.l.* lineage, suggesting a single genomic entity. Group B’s *Iberian clade* incorporated expected species (*Q. canariensis, Q. faginea*, *Q. lusitanica* and *Q. pyrenaica*). Unlike in the maximum likelihood (concatenation) analysis (Fig. 2), *Q. canariensis* samples separated into two separate clades in the SVDquartets analysis (Fig. S1). Samples from the Iberian SW (Monchique (Portugal) and Algeciras (Spain)) remained in the *Iberian clade*, while samples from NE Spain (Catalunya) clustered within the *pubescens clade*, which contains all *Q. pubescens* samples with other Southern European origins.

### Population structure analysis

Samples were subsequently resolved using Principal Component Analysis (PCA), which sorted them into phylogenetic Groups A and B according to PC1 (7.4% of variance), while *Quercus lusitanica* separated along PC2 (2.9% of variance) (Fig. 3A). In a PCA of Group A species (Fig. 3B), PC1 separated European *Q. robur* and *Q. petraea s.l.*, from the Iberian representatives of *Q. robur s.l.* Within the latter assemblage, samples initially assigned to *Q. orocantabrica* did not separate from those assigned to *Q. broteroana*, whereas *Q. estremadurensis* remained separate and partially displaced towards *Q. petraea s.l.*. The PCA of Group B species (Fig. 3C) again separated *Q. lusitanica* from the other species, along PC1 (4.3% of variance). PC2 (2.7% of variance) reflected biogeography, gradually resolving Near East *Q. boissieri, Q. kotschyana,* the Anatolian and Caucasian *Q. vulcanica* and *Q. macranthera,* followed by *Q. pubescens*, and finally all Iberian Group B species. To complement this information, we conducted a Neighbor-net analysis that showed some reticulation at the bases of the clades, although within-species replicate samples largely clustered together (Fig. S2). This analysis recovered phylogenetic groups A and B, except for *Q. estremadurensis*, reflecting its low split length. Notably, in group A, the European *Q. robur* and the Northwestern Iberian lineage of samples initially labeled as *Q. broteroana*/*Q. orocantabrica*, were differentiated from each other and from all remaining species. Also of significance, species were broadly resolved between Iberian and non-Iberian provenances, regardless of their Group A or B attribution. For instance, inside the *Q. petraea s.l.* group, a splitting was observed between the Iberian *Q. pauciradiata* and the remaining European *Q. petraea*.

**Fig. 3.**
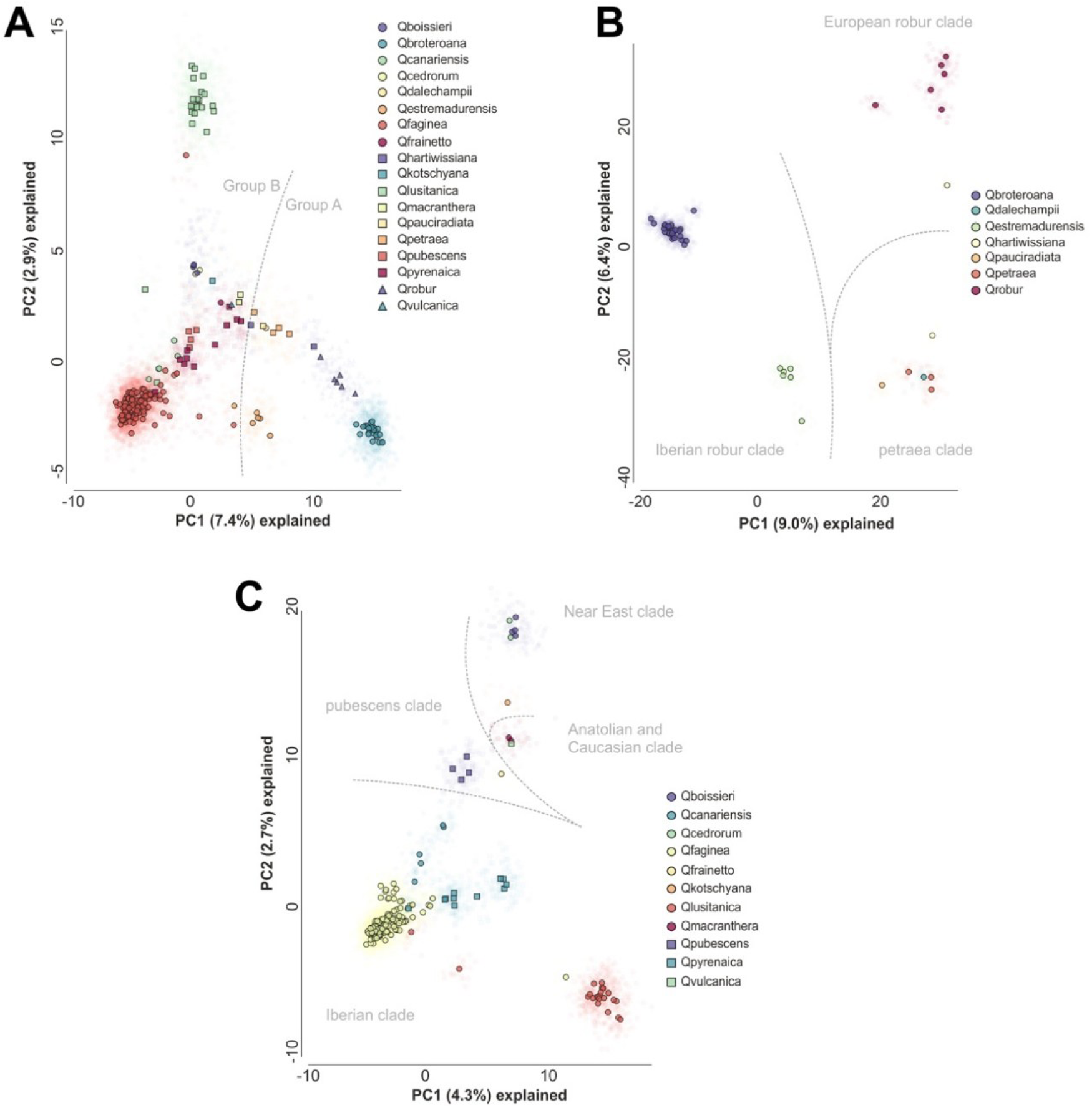
Principal component analysis of Eurasian white oaks. (A) Global overview with all species; dashed grey line delimits Groups A and B. (B) Group A species; dashed grey line delimits the presently established Group A clades (see Fig. 1); (C) Group B species; dashed grey line delimits the presently established Group B clades (see Fig. 1).

### Evaluation of nothotaxa relationships

Subsequently, we extended our analysis to populations of hybrids (known nothotaxa) using PCA, while also incorporating ancestrality testing into our insights. The first analysis consisted of a birds-eye view of all sampled populations following incorporation of hybrids (Fig. 4A). Here, we also observed the segregation between Groups A and B reflected onto PC1 (6.0% of variance). PC2 (2.4% of variance) translated a Mediterranean East-to-West gradient. Our dataset did not incorporate any tentative hybrids within Group A species. This was reflected in the ensuing analysis for patterns of ancestry within group A, using STRUCTURE (Figs. 4B and S3). The best fit (K=3) resolved clusters for *Q. robur*, *Q. petraea s.l.*, and a single Iberian cluster comprising samples initially assigned to *Q. broteroana* and *Q. orocantabrica*. *Quercus estremadurensis* showed strong evidence of shared ancestry between *Q. petraea s.l.* and the Iberian *Q. broteroana–Q. orocantabrica* lineage.

**Fig. 4.**
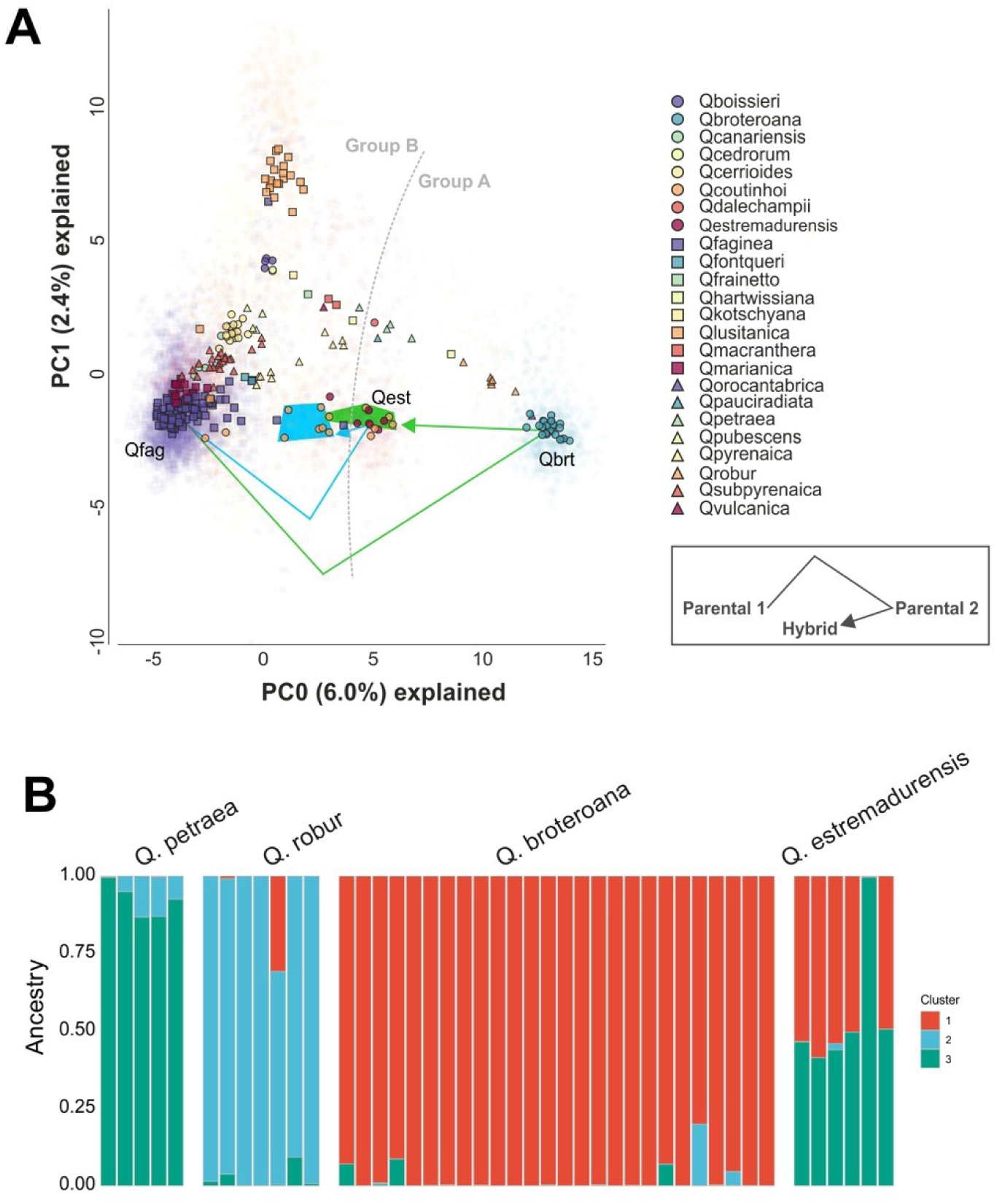
Population structure analysis of Eurasian white oaks. (A) Principal component analysis global overview, with all species; dashed grey line delimits Groups A and B. (B) STRUCTURE analysis of ancestry proportions of Group A samples (K=3).

Group B contained several tentative intra-group hybrids and was subjected to a group-specific PCA (Fig. 5A). Results consistently evidenced the studied nothotaxa in intermediate positions between putative parents. The first segregation level (PC1; 3.3% of variance) resolved *Q. lusitanica* samples, followed by the eastern distributed species (*Q. boisseri, Q. frainetto, Q. kotshyana, Q. macranthera,* and *Q. vulcanica*) and *Q. pyrenaica.* Meanwhile, PC2 (2.7% of variance) reflected multiple hybrids and parent species. Top-to-bottom, segregation evidenced a gradient between *Q. pubescens,* followed by *Q. ×cerrioides* and the Catalonian samples of *Q. canariensis*. This hybrid swarm was followed by *Q. ×subpyrenaica* intermediate with *Q. faginea*. Finally, samples of *Q. ×marianica* were also positioned between *Q. canariensis* and *Q. faginea*. Ancestry analysis preferably assigned six clusters that cohesively resolved the five species present in our dataset, plus sub-structuring of the extensive *Q. faginea* sampling (Figs. 5B and S4). Of significance, four populations of hybrids matched expected ancestry components for their assumed parental species. Neighbor-net analysis of the complete hybrid dataset mirrored previous results, showcasing the relationships between the main species and putative parental relationships (Fig. S5). A summary of parent/hybrid lineages is depicted in Fig. 5C.

**Fig. 5.**
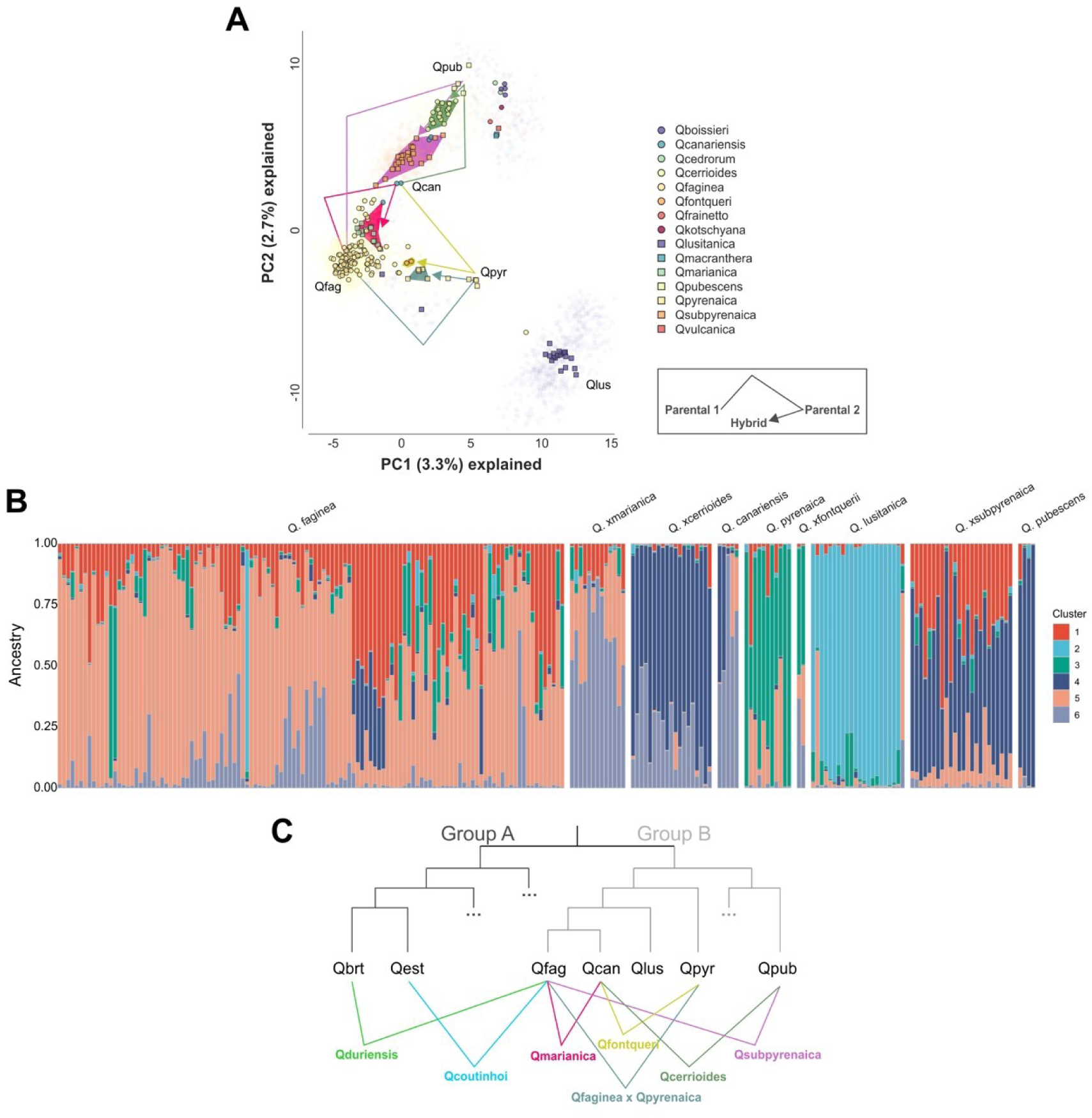
Population structure analysis of Eurasian white oaks. (A) Principal component analysis of Group B species; dashed grey line delimits the presently established Group B clades. (B) STRUCTURE analysis of ancestry proportions of Group B samples (K=6). (C) Schematic representation of the relations between hybrids and parent species. Qbrt, *Q. broteroana*; Qcan, *Q. canariensis*; Qest, *Q. estremadurensis*; Qfag, *Q. faginea*; Qlus, *Q. lusitanica*; Qpub, *Q. pubescens*; Qpyr, *Q. pyrenaica*.

Within our cohort, the intergroup A/B hybrids represent a special case. A combination of literature and biogeographic evidence led to grouping two sets of hybrids under the name *Q. ×coutinhoi*. They share the same Group B parent species (i.e., *Q. faginea*) but differ in the Group A parent species (the Iberian *robur* clade members *Q. estremadurensis* and *Q. broteroana*). We ran PCA analysis incorporating two tentative *Q. ×coutinhoi*. One, sympatric with *Q. estremadurensis*, was visible halfway between *Q. estremadurensis* and *Q. faginea* (Figs. 4A and S6A). A second, sympatric with *Q. broteroana*, was visible halfway between *Q. broteroana and Q. faginea*, albeit superimposing with *Q. estremadurensis* (Figs. 4A and S6B). A new PCA that removed the confounding effect of remaining Group B members was able to separate the latter *Q. ×coutinhoi* from *Q. estremadurensis* (Fig. 6). By resolving both *Q. ×coutinhoi* samples at the population level, we were able to observe a tentative origin from two separate provenances (*Q. broteroana × Q. faginea* and *Q. estremadurensis* × *Q. faginea*). Results support the previous nomenclatural attribution and taxonomic rearrangement provided by Vasconcellos and Franco (1954) of *Q. ×duriensis,* which we advocate for the hybrid between *Q. broteroana* and *Q. faginea* (Fig. 5C; Note S1). Our results reinstate the need for in-depth molecular characterization in support of sophisticated taxa and nothotaxa attribution.

**Fig. 6.**
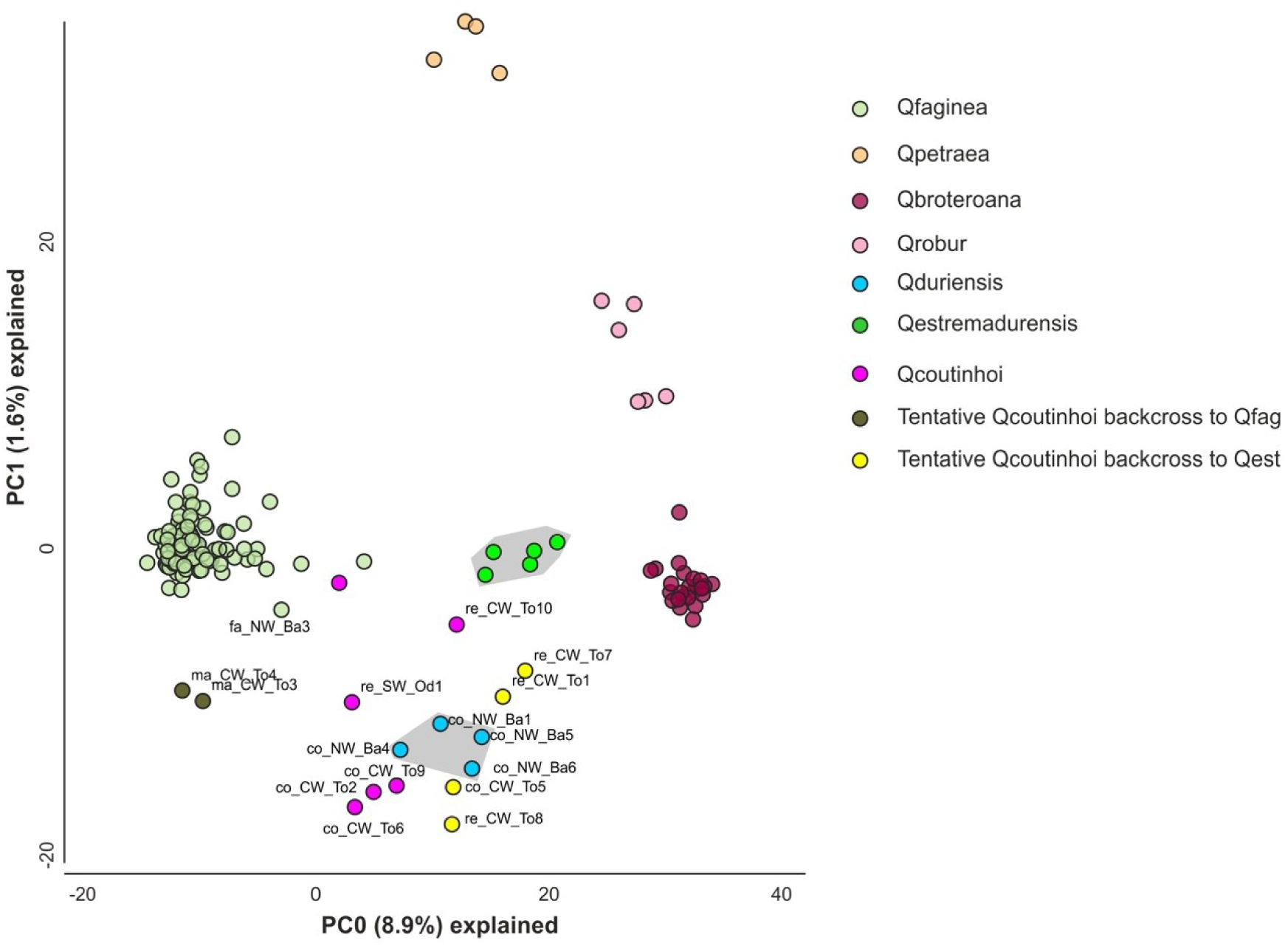
Principal component analysis addressing hybridization in Iberian white oaks, removing the confounding effect of Group B members in order to resolve *Q. ×coutinhoi* from *Q. estremadurensis* in the bi-dimensional space (grey highlighted areas).

## Discussion

### The European white oaks reflect two new major groups

Radiation within Eurasian *Roburoid* (Sect. *Quercus*) oaks has been a longstanding topic of interest for plant evolution. Here, the characterization of a large set of Iberian samples led to the novel observation of two well defined clades (Groups A and B), which reveal two new subsections inside Section *Quercus*. This hypothesis contradicts the previous assignment of four subclades, all of which carried dry-adapted (*Galliferae*) species and were suggestive of convergent evolution towards adaptation (Hipp *et al*., 2020). The imperfect identification of *Galliferae* representatives (*Q. canariensis* and *Q. lusitanica*) (Hipp *et al*., 2020) possibly contributed to this misassignment. In the present report, the phylogenetic analysis of the Iberian white oaks revealed deep intraspecific interconnection and solid geographic structuring in samples from both groups (A and B), which were ascribed to former taxonomic groups (sections), but with different and newly solved circumscriptions. Group A incorporates the temperate true roburoid species (*Q. robur s.l.* and *Q. petraea s.l.*). Group B species include mostly circummediterranean marcescent oaks, which occur predominantly in the transitional areas towards the Mediterranean climate (De Dios *et al*., 2009; Mauri *et al*., 2016; Vilches *et al*., 2016; Dorado-Liñán *et al*., 2017; Gil-Pelegrín *et al*., 2017; Kavgacı *et al*., 2021). Their distributions correlate with an increase of annual and summer precipitation, which confers a submediterranean character to those forest ecotones, between fully deciduous and evergreen sclerophyll species (García-Mijangos *et al*., 2015; Hipp *et al*., 2020; Vila-Viçosa *et al*., 2020b).

### Group A members highlight the uniqueness of Iberian oak diversity

Our findings highlight three taxonomic novelties in the Iberian white oaks Clade A, which, in the past, were largely overlooked. First, the Iberian *Quercus pauciradiata* is noticeably divergent from its European *Q. petraea s.l.* counterparts, as suggested by branch length (Figs. 2 and S1) and separation in the PCA (Fig. 3B). This result is supported by *Q. pauciradiata*’s described morphological variation, with an exclusive and different type of fasciculate indumentum (Penas *et al*., 1997) when compared to the remaining *Q. petraea s.l.*. It resembles the example of another Southern European *Q. petraea s.l.* oak, the Italian *Q. dalechampii,* which has been taxonomically ambiguous due, at least in part, to morphological plasticity (Di Pietro *et al*., 2020; Proietti *et al*., 2021).

Second, our results support the existence of two Iberian *Q. robur s.l.* lineages, corresponding to either *Q. estremadurensis*, or the Northwestern Iberian lineage represented in our sampling by material initially assigned to *Q. broteroana* and Q*. orocantabrica*. Both lineages are consistently differentiated from the remaining European *Q. robur s.l.*, supporting the distinctiveness of the Iberian pedunculate-oaks, which may reflect legacies of past population dynamics associated with glacial refugia (Olalde *et al*., 2002). Samples initially assigned to *Q. orocantabrica* were nested with *Q. broteroana*, and neither the PCA nor the population-structure analyses support their recognition as independent evolutionary lineages. Although *Q. orocantabrica* was previously described as a distinct species (Rivas-Martínez *et al*., 2002), we advocate that both names are better treated as referring to a single taxonomic entity, for which formal nomenclatural assessment should be implemented (Note S1).

Third, the phylogeny, PCA and ancestry analyses corroborate an interesting pattern around *Q. estremadurensis*, suggestive of a shared ancestry with *Q. petraea s.l.* and *Q robur s.l.* inside Group A. Morphologically, this is reflected in *Q. estremadurensis*’ features such as a larger petiole and rhomboidal leaf shaping (Schwarz, 1935; Schwarz, 1936a; Vicioso, 1950; Vila-Viçosa *et al*., 2014; Vázquez *et al*., 2018). It remains to be established whether this reflects a past hybridization event between lineages, or the presence of ancestor footprint in *Q. estremadurensis* populations, in which case the Western IP may have acted as a putative center of origin for the temperate oaks (Group A). These results place the Iberian Group A taxa within a broader southern European context of roburoid and petraeoid diversity, marked by historical taxonomic complexity, intermediate morphologies, and biogeographic ambiguity (Note S2).

### Group B species reflect an East-to-West geographic gradient

Group B species segregate between the *Iberian* and *pubescens* clades, the former encompassing Iberian samples but reflecting a broader geographical distribution of species across the Western Mediterranean. The latter clustered with Anatolian, Caucasian and Near East species (Fig. 2 and 3C). This observation broadly reflects an East-to-West biogeographical structuring across the Mediterranean basin. *Quercus canariensis* is an interesting case of this biogeographic continuum. Our analysis revealed that a subset of Catalonian samples was geographically related to *Q. pubescens* and *Q. ×cerrioides* (which we interpret as a hybrid swarm of *Q. canariensis* x *Q. pubescens*) (Figs. 5B, S1 and S5). Results indicate that Catalonian populations of *Q. canariensis* display high levels of admixture with southern France and Italian *Q. pubescens*. Most importantly, the molecular evidence for this East-to-West continuum seems to contradict previous assignment of Western Mediterranean *Q. faginea, Q. canariensis*, *Q. lusitanica* and the Near Eastern *Q. boissieri*, into a single Section *Galliferae* (Boissier, 1853; Gürke, 1897; Tschan & Denk, 2012). Our molecular evidence supports a scenario of geographic structuring, compatible with vicariance, between *Q. canariensis* and *Q. boissieri* at opposite margins of the Mediterranean distribution (Figs. S1 and S5), with *Q. pubescens* occupying a broader, geographically intermediate position. Similarly, four species with morphological similarities (the Iberian *Q. pyrenaica,* the Italo-Hungarian *Q. frainetto*, the Anatolian *Q. vulcanica* and the Lebanese *Q. kotschyana*), all traditionally assigned to former Series *Confertae* (Section Dascia) (Schwarz, 1936a), appear in our data as phylogenetically segregated. Such evidence supports an additional case of either convergent evolution, phenotypic plasticity, or common ancestry with posterior gene flow with local species, deserving deeper population genetics and phylogenetic surveys to infer on past demographic history across southern Europe.

Another interesting finding relates to *Q. lusitanica*, a species consistently assigned to Group B (Figs. 2 and S1). In PCAs, *Q. lusitanica* was responsible for much of the observed variance, whether analysis referred to Group B species or the global oak analysis (Fig. 3A,C, Fig. 4A, Fig. 5A). Unlike remaining European white oaks, *Q. lusitanica* is a shrub. It has been morphologically and historically associated with *Q. faginea*, resulting in ancient nomenclatural entanglement with this species (Coutinho, 1888; Sampaio, 1910; Vicioso, 1950; Vázquez *et al*., 2018). This shrub alternates between marcescent and evergreen leaf behavior. Ecologically, it is limited to siliceous bedrocks, requires moderate annual precipitation, and is highly susceptible to winter cold, making its distribution restricted to the Atlantic front of the IP and North Africa (Capelo *et al*., 2002; Vila-Viçosa *et al*., 2020a; Vila-Viçosa *et al*., 2020b). This ecological specificity, coupled with it being a geoxylic shrub, may have led to genetic divergence by selection. Alternatively, neutral genetic differentiation may have resulted from genetic bottlenecks associated with severe dry and cold periods or adaptation to cycles of recurrent fires, in stages of degradation of the Mediterranean forest into shrubland and grassland mosaics observed in the western Iberian Peninsula in the past, especially during hot periods (Daniau *et al*., 2007; Connor *et al*., 2012; Camuera *et al*., 2019; Genet *et al*., 2021). This pattern should be interpreted within the broader natural-history context of Mediterranean and submediterranean oak differentiation, including the role of transitional ecotones in shaping regional diversity (Note S2).

### The Iberian white oaks reflect a legacy of pervasive hybridization

Our analysis demonstrated intermediate positioning of numerous hybrids between their putative parents, validating classic nothospecies (Schwarz, 1936a; Vicioso, 1950; Villar, 1958; Rivas-Martínez & Saénz, 1991; Schwarz, 1993; Vila-Viçosa *et al*., 2014). More specifically, using a combination of morphological identification, literature review (Note S1) and molecular inference, we were able to assign a large set of Group B intra-hybrids, and two (albeit complex) Group A-B inter-hybrids. The latter showed a gradient with putative hybrids positioned intermediately among parents, going as far as to suggest cases of backcrosses (Fig. 6, and S6). Results respectively associate *Q. ×duriensis* and *Q. ×coutinhoi* with known hybrid swarms that occur either in the Northern Douro Basin (for the contact between *Q. faginea* and *Q. broteroana*), or the Center and Southwest Portugal (for the contact between *Q. faginea* and *Q. estremadurensis*) (Vila-Viçosa *et al*., 2020b).

Group B concentrated the strongest evidence of syngameon dynamics, with a higher number of geographically widespread hybrid swarms involving four species and several intermediate nothotaxa, namely *Q. ×cerrioides, Q. ×subpyrenaica*, *Q. ×marianica* and *Q. faginea × Q. pyrenaica* (Fig. 5B). Figs. 5B and S5 highlight the Northeastern Iberia hybrid swarm composed of *Q. canariensis*, *Q. pubescens* and their hybrid *Q. ×cerrioides*, as well as the intermediate behaviour of hybrid swarms in relation to parent samples. We highlight *Q. faginea* as one of the species most prone to hybridize with multiple Iberian oaks (Fig. 5B). This might be due to the ecological plasticity of *Q. faginea,* as the most dry-adapted species of all Iberian white oaks, spanning across typical Mediterranean areas but also capable of contacting with temperate and continental areas (Kouba *et al*., 2011; Vila-Viçosa *et al*., 2012; Amigo *et al*., 2017; Vila-Viçosa *et al*., 2020b).

### The Iberian Peninsula syngameon as a hotspot for oak conservation

A major output of the present work is the resolution of the phylogenetic backbone and species validation for forest dominant trees, in an important area (the IP) placed between two major biogeographic subregions (Eurosiberian and Mediterranean). First, by refining the taxonomy inside the *Q robur s.l.* group, we showcase *Q. estremadurensis* and *Q. broteroana* as taxa that should be considered as species due to the considerable phylogenetic divergence with European pedunculate oaks (*Q. robur* L. *senso strictu).* Earlier, we also highlighted a potential Iberian origin for Group A oaks, based on molecular and morphological data coming from *Q. estremadurensis*. *Q. estremadurensis* is distributed across Western IP in very small patches of scattered individuals among *Q. faginea* forests (Vila-Viçosa *et al*., 2020b) where, frequently, the presence of *Q. ×coutinhoi* is dominant over *Q. estremadurensis*. Thus, this species is deserving of major conservation efforts and inclusion in the *Species of Concern and Genetic Research Focus* list (Backs & Ashley, 2021). Second, the lineage initially assigned to *Q. broteroana* and *Q. orocantabrica*, appears to merit recognition as a distinct species-level entity within *Q. robur s.l..* Further population genomic studies are needed to assess geographic substructure within this lineage, particularly across northwestern Iberian and Cantabrian populations, and to clarify the nomenclatural treatment required by the priority rule (Turland *et al*., 2025). Similarly, *Q. pauciradiata*, which belongs to the *Q. petraea s.l.* in our analysis, also deserves further studies alongside *Q. huguetiana,* an Iberian petraeoid oak appearing as a sister species to *Q. petraea* (Papini et al., 2011). Finally, the validation of several nothotaxa, based on consistent population structure evidence, urges for novel conservation efforts in sensitive areas such as the Catalonian sub littoral mountains (*Q. ×cerrioides*), the Pre-Pyrenean (*Q. ×subpyrenaica*), and the ongoing introgression of both *Q. broteroana* and *Q. estremadurensis* with *Q. faginea* related with *Q. ×duriensis* and *Q. ×coutinhoi* in Western Iberia. Much like with American white oaks, this peripheral natural hybrid zones emerge as notable case studies to address adaptation and adaptive introgression of genomic variation (Suarez-Gonzalez *et al*., 2016; Holliday *et al*., 2017; Ribicoff *et al*., 2025). Our conservation concerns are even more pressing in the face of the fast pace of climate change, where bioclimatic and ecological adaptation will undoubtedly play a significant role in species migration, survival rates, and isolation, thus shaping sympatry patterns and exposure to introgression with other oaks.

## Concluding remarks

The present analysis contributed to the resolution of several pending taxonomic questions, summarized into three main findings. First, Iberian oak species were successfully readdressed, enlarging the current understanding of the phylogeography of the Eurasian white oaks. Second, our infrasectional analysis resulted in the important identification of two subsections (Groups A and B) in the western Paleartic white oaks, tackling a key challenge in oak evolution – resolving oak taxonomy below the section level (Denk *et al*., 2017). Third, the detection of hybrid swarms confirms the extensive gene flow among oak populations, despite the maintenance of parental lineages and integrative interbreeding evolutionary units (Suarez-Gonzalez *et al*., 2018). Ultimately, our approach improves on the knowledge obtained at the micro- and macroevolutionary levels in the European white oaks, after the development of oak genomics (Plomion *et al*., 2016; Sork *et al*., 2016; Ramos *et al*., 2018; Leroy *et al*., 2020b; Zhou *et al*., 2021; Fu *et al*., 2022; Liang *et al*., 2022; Sork *et al*., 2022). It solidifies the Iberian Peninsula as a hotspot for white oak diversity, urges the development of novel conservation strategies, and emphasizes the role of Southern European Peninsulas as reservoirs of oak biological diversity.

## Acknowledgments

We acknowledge the insightful discussions with Professor Guido Grimm on white oak evolution, which greatly enriched this paper. The authors acknowledge all collaborators, friends, and oak enthusiasts who helped survey Iberian oak populations across the range. CVV received support from the Portuguese Ministry of Education and Science and the European Social Fund, through the Portuguese Foundation of Science and Technology (FCT) (contracts PD/BD/52607/2014 and project UID/50027/2025). This article expands upon material previously presented in the author CVV’s PhD thesis.

## Author contributions

CVV, RAC, and HA conceived and designed the study. CVV and FMV collected, identified, and curated samples. CVV, RAC and HA generated and analyzed phylogenetic and genomic data, with support from ABP and AH. CVV, RAC, CG and HA drafted the manuscript. AH supervised and provided key insights into the final version of the manuscript, alongside CC and RA. All authors wrote and edited the manuscript.

