## Supplementary Material for "The Iberian white-oak syngameon as a legacy of introgression in southern Europe"

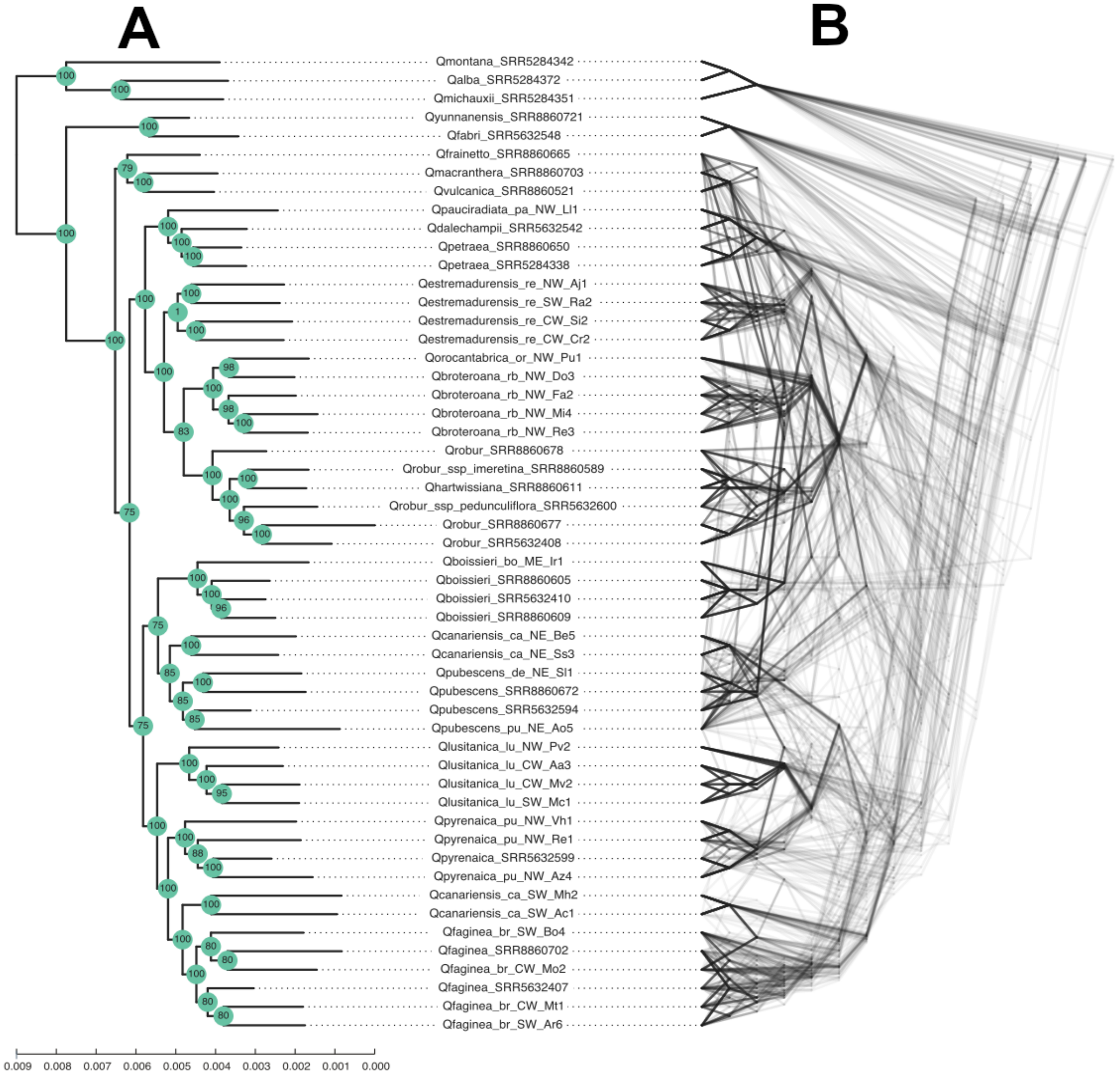


**Fig. S1** Phylogeny of Iberian white oaks Using three American oaks (*Q. alba, Q. michauxii, Q. montana*) and two Asian oaks (*Q. fabri* and *Q. yunnanensis*) as outgroups. Maximum likelihood (ML) analyses on the concatenated matrix (A) and species tree inference based on the SVDQuartets algorithm (TETRAD) (B).

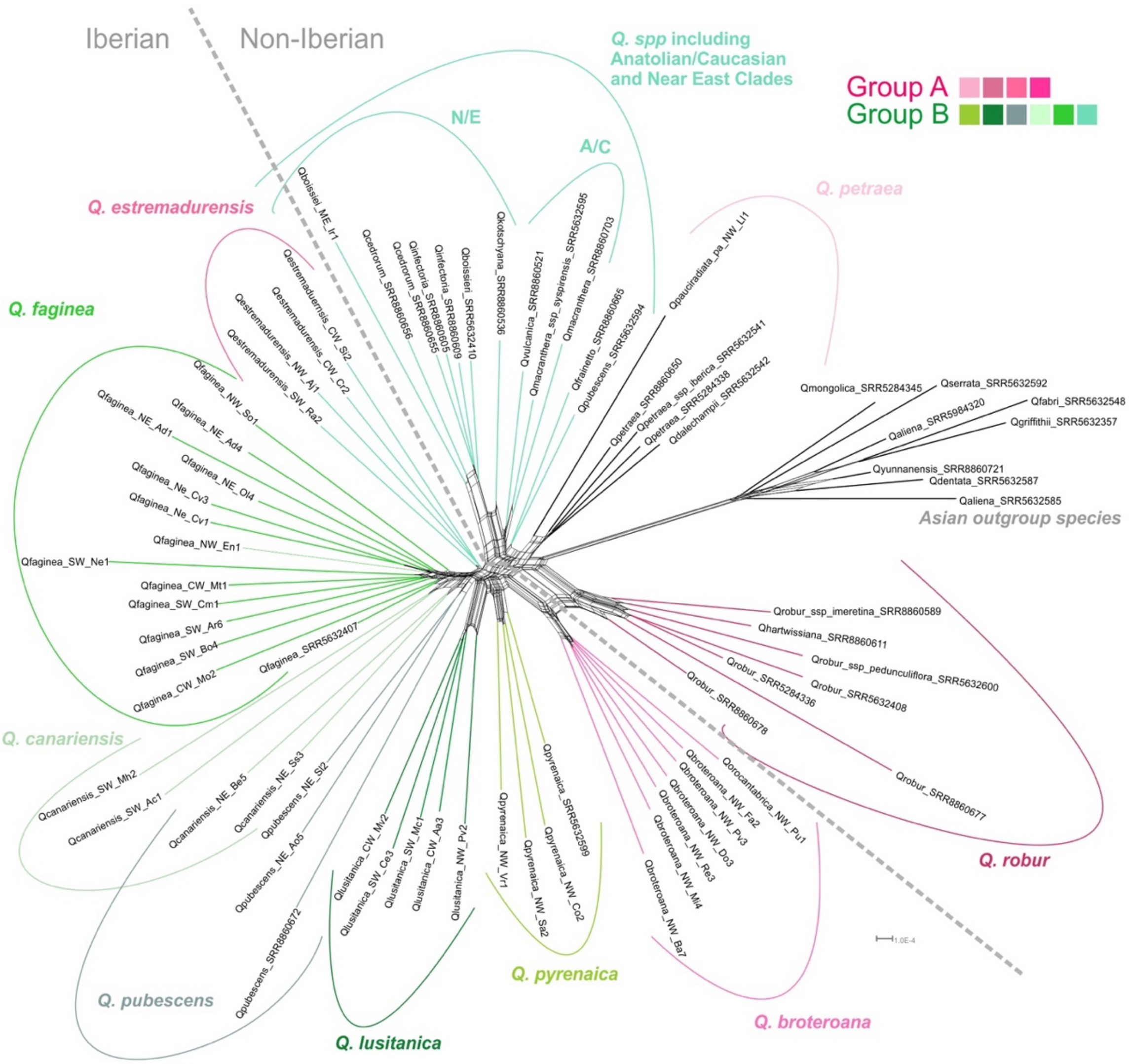


**Fig. S2** Phylogenetic network computed using the neighbor-net method based on pairwise maximum likelihood distances and focusing on a species-level analysis of Iberian white oaks and related taxa. Colored lines group phylogenetically cohesive species samples from Group A (purples), Group B (greens), and the outgroup Asian species (light grey). Dashed grey lines represent a large biogeographic split delimiting Iberian from non-Iberian samples.

Outgroup Asian species (separated to the right), clustered closer to Group A species. In Group A, both *Q. robur s.l.* and *Q. petraea s.l.* recovered Iberian taxa (*Q. broteroana* and *Q. pauciradiata*). Group B species incorporated a subset of European and Euroasiatic species, where one sample of *Q. pubescens* grouped with *Q. frainetto*, followed by Caucasian (Q*. macranthera*) and Anatolian (Q. *vulcanica*) samples. Curiously, the Near East *Q. kotschyana* shared some splitting between *Q. vulcanica* and *Q. boissieri*, complementing the Near East clade. The Iberian set of Group B samples resolved the main species (*Q. faginea*, *Q. canariensis*, *Q. lusitanica,* and *Q. pyrenaica*), while reflecting the transitional gradient of Catalonian *Q. canariensis* samples towards *Q. pubescens*.

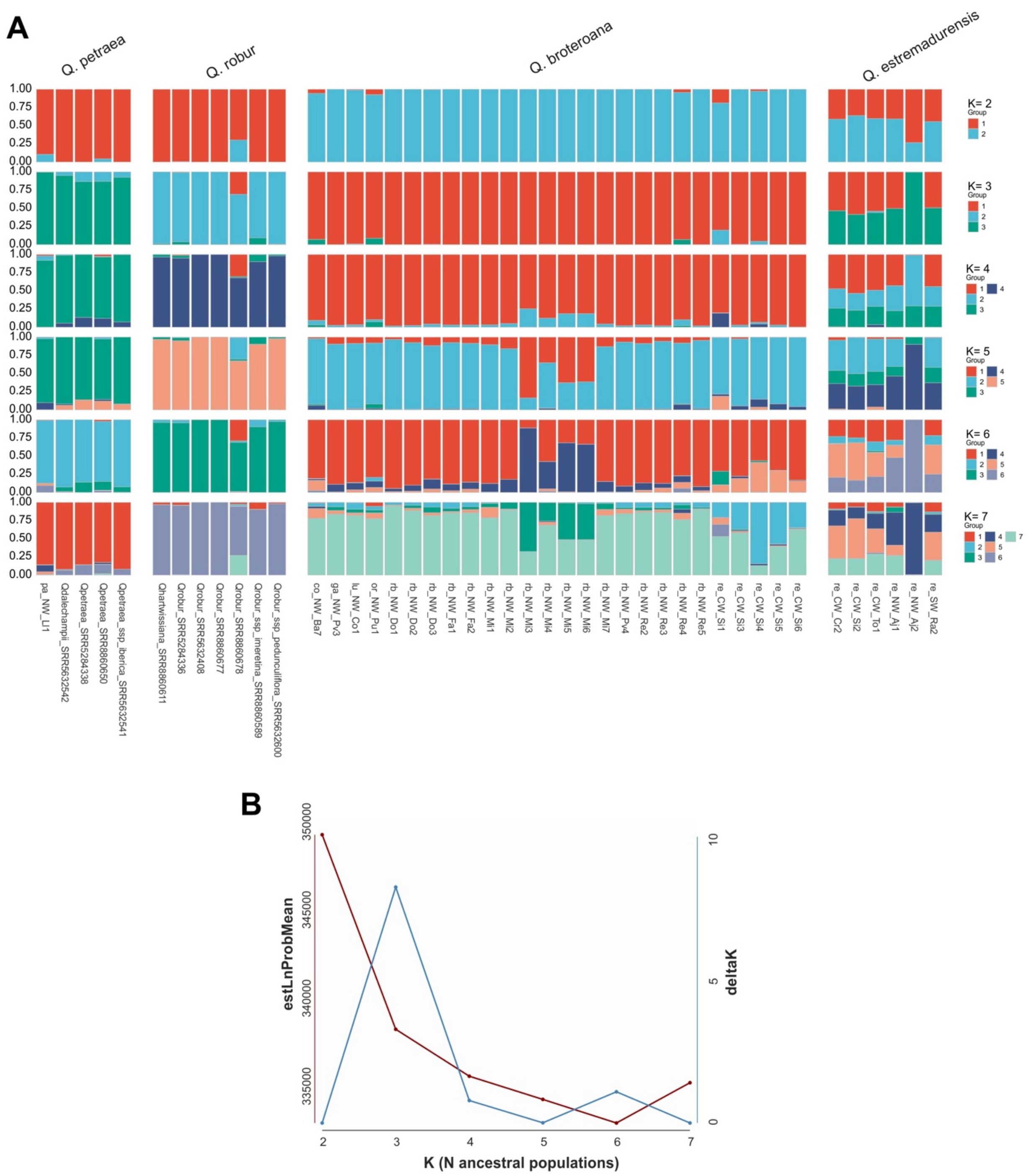

**Fig. S3** STRUCTURE analysis of Group A Eurasian white oaks. (A) STRUCTURE results for Group A species (*Q. petraea*, *Q. robur*, *Q. broteroana* and *Q. estremadurensis*), with K values ranging from 2 to 7. Each vertical bar represents an individual, and colors indicate the proportion of ancestry assigned to each inferred ancestral population. (B) Evaluation of the most likely number of ancestral populations (K), showing the mean estimated log probability of the data (estLnProbMean) and ΔK values.

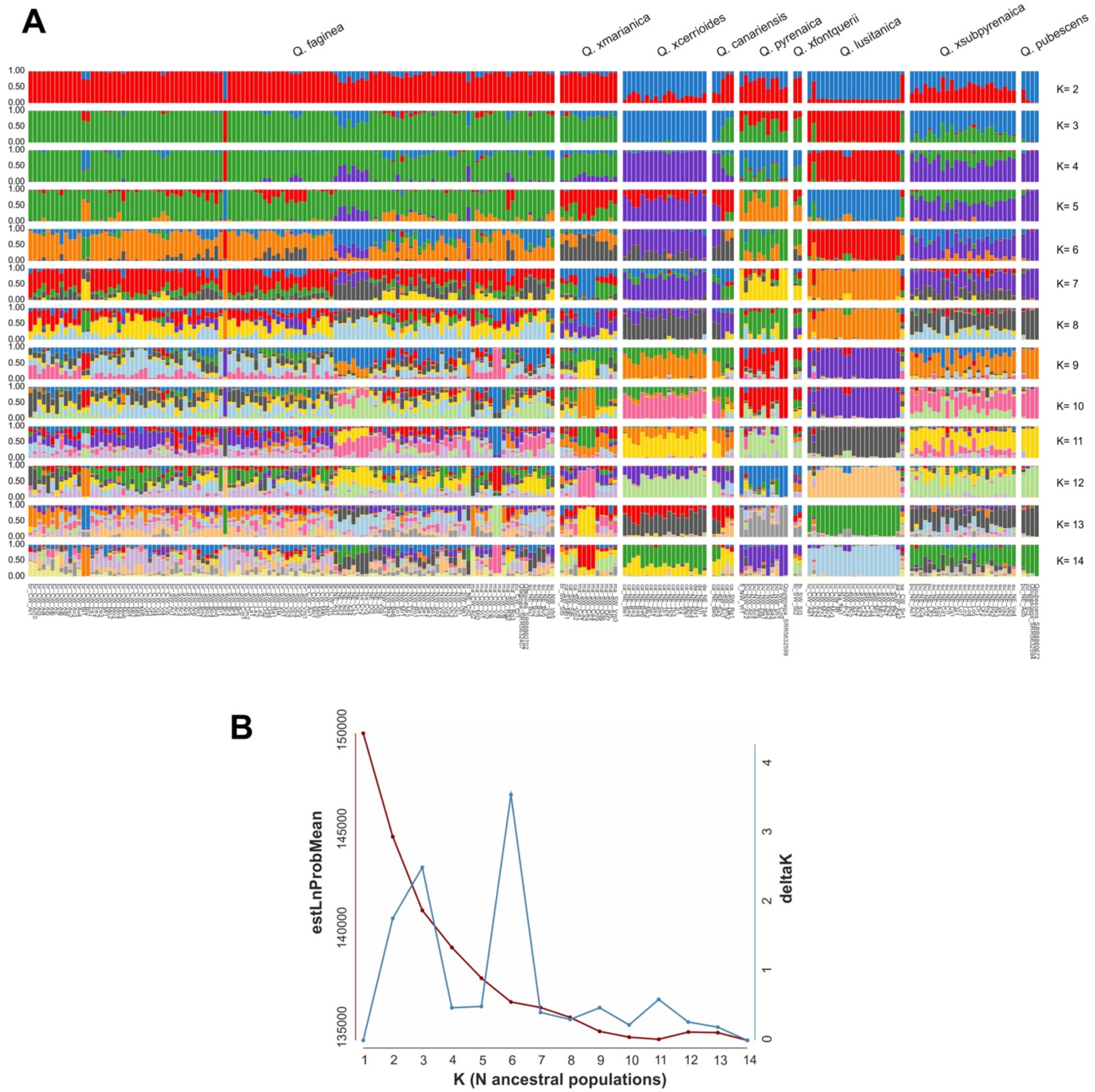


**Fig. S4** STRUCTURE analysis of Group B Eurasian white oaks. (A) STRUCTURE results for Group B species (including *Q. faginea*, *Q. canariensis*, *Q. pyrenaica*, *Q. lusitanica*, *Q. pubescens*, and related nothtotaxa, with K values ranging from 2 to 14. Each individual is represented by a vertical bar, and colors correspond to the estimated ancestry proportions for each inferred ancestral population. (B) Evaluation of the most likely number of ancestral populations (K), showing the mean estimated log probability of the data (estLnProbMean) and ΔK values.

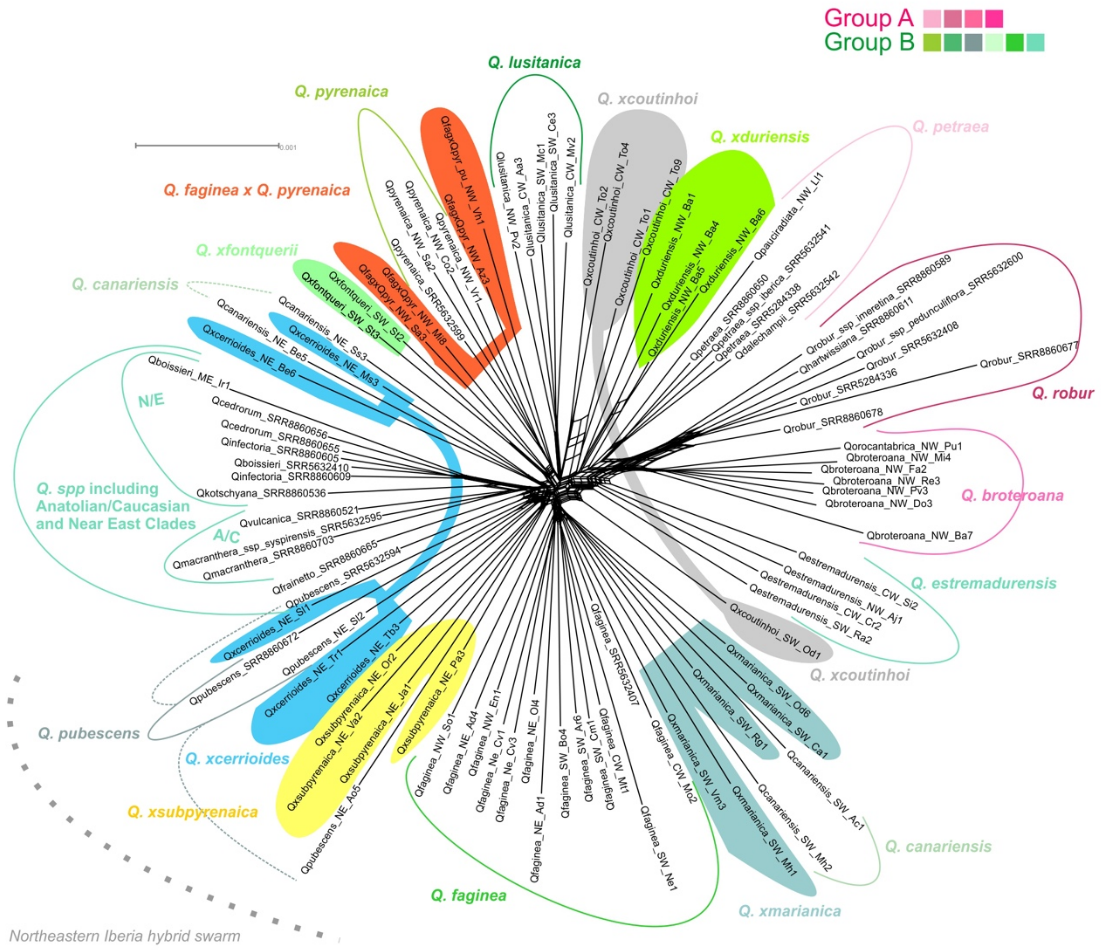


**Fig. S5** Neighbor-net analysis focusing on Iberian white oaks, including their hybrids. Colored lines group phylogenetically cohesive species samples from Group A (purples) and Group B (greens). Colored fill areas group cohesive hybrid samples.

We observed an expected confounding effect resulting from the incorporation of hybrids, as shown by the relative splitting among parents. The four hybrids formed with *Q. faginea* displayed either intermediate positioning among parents (*Q. ×marianica* and *Q. ×subpyrenaica*) or positioning close to one parent (*Q. faginea × Q. pyrenaica* and Q. *×coutinhoi*). We observed a Northeastern Iberia hybrid swarm, which incorporated the adequately positioned *Q. ×subpyrenaica*, and Catalonian *Q. ×cerrioides* samples, the latter being equally distributed among the parents *Q. canariensis* and *Q. pubescens*. Meanwhile, *Q. ×fontquerii* (*Q. canariensis × Q. pyrenaica*) correctly adopted an intermediate position between parents.

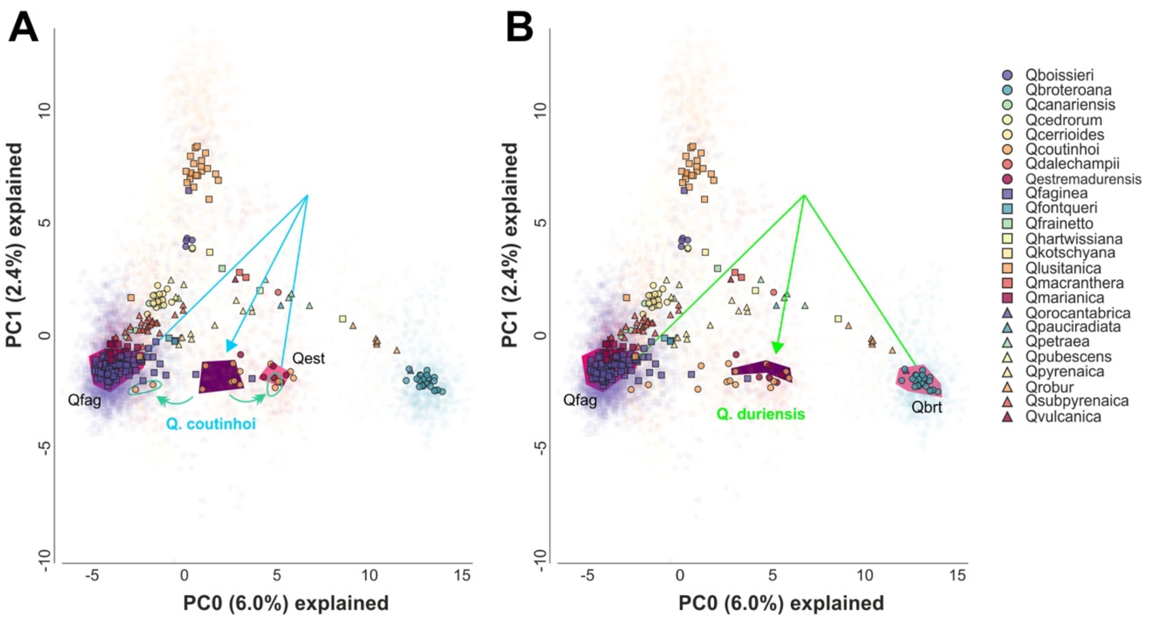


**Fig. S6** Principal component analysis addressing hybridization in Iberian white oaks. (A) PCA featuring all taxa and hybrids, highlighting samples that reflect the hybrid *Q. ×coutinhoi* and tentative backcrosses (green arrows). (B) PCA featuring all taxa and hybrids, highlighting samples that reflect the hybrid *Q. ×duriensis*.

**Method S1** Characterization of the study area.

The Iberian Peninsula is located in the southwest of the European continent and the extreme west of the Mediterranean Basin (between latitudes -9.55° W to 3.35°E and latitudes 35.87° N to 43.80° N). The IP holds a highly diverse geological history, with a geographic location that places it under the influence of two strong climatic determinants: a wet Temperate Atlantic climate in the Northwest, and a dry Mediterranean climate in the Southeast (Loidi, 2017; Rivas-Martínez *et al.*, 2017). Its biogeographic positioning and orographic characteristics turn it into a hotspot for European phytocoenotic diversity. The Iberian total flora displays almost 30% of endemicity rate (including 21 genera and 13 phylogeographic refugia) (Aedo *et al.*, 2017; Médail & Baumel, 2018), and it specifically hosts circa half of the European oak species (Schwarz, 1993).

**Note S1** Information regarding the focal taxa addressed in the present report.

**Section *Quercus*** Denk *et al.* (2017)

**Subsection *Robur*** Schwarz (1936a)

***Quercus estremadurensis*** O. Schwarz *(=Q. robur* subsp. *estremadurensis* (O.Schwarz) A. Camus)

**General information:** Taxon earlier described by Schwarz (1935) as an independent species of *Q. robur* L.. Together with *Quercus hartwissiana* Steven, both species present common characteristics that led the author to create a Series "Primitivae", inside Section *Robur* with characters related with cup scales and leaf development, morphology and venation (Schwarz, 1936b). Both develop large petioles, and petraeoid leaves. In the original protologue, the author already refers problems of introgression with sympatric *Quercus robur*, assuming that all the plants that he studied are already severely hybridized with the "European" *Quercus robur*. This taxon is considered to be relictual and distributes across western Iberian Peninsula, especially in thermophilic areas and river basins. Related vouchers from Northern Africa (Paris Herbarium) can also be addressed to this taxon (Vila-Viçosa *et al.*, 2014; Vázquez *et al.*, 2018).

**Differential morphological traits:** Regular and rhomboidal leaf structure, relative higher number of acute and equal lobes and often absence of sinual nerves, the higher number of scales, which are large at the base of the cup and becoming smaller towards the apex. It tends to develop large petioles (up to 1 cm) and large peduncles (>10 cm) (Schwarz, 1935; Vila-Viçosa *et al.*, 2014; Vázquez *et al.*, 2018).

***Quercus broteroana* / *Quercus orocantabrica* complex** (=*Q. robur* subsp. *broteroana* O.Schwarz; *Q. orocantabrica* Rivas Mart., Penas, T.E.Díaz & Llamas)

**General information:** The Northwestern Iberian pedunculate-oak lineage has historically been treated under different names and taxonomic ranks within the broad *Q. robur s.l.* complex. *Quercus robur* subsp. *broteroana* was described by Schwarz shortly after *Q. estremadurensis*, as an Iberian subspecies interpreted by the author as part of the putative introgressive gradient between *Q. estremadurensis* and European *Q. robur* L.. Schwarz emphasized leaf, cupule, petiole and peduncle dimensions as diagnostic features, together with an essentially Northwestern Iberian distribution and a biogeographic frontier with

European *Q. robur* in northern León, Spain (Schwarz, 1937). Later, in Flora Europaea, Schwarz subsumed both *Q. estremadurensis* and *Q. robur* subsp. *broteroana* within a broader *Q. robur* concept, after observing partly convergent character states among European *Q. robur s.l.* material Schwarz (1964). Nevertheless, his observations on Iberian pedunculate oaks remain relevant, particularly because the characters he highlighted occur at higher frequency in Northwestern Iberian material, as also supported by the genomic structure recovered in the present study.

*Quercus orocantabrica* was subsequently described as a relict roburoid oak endemic to the Cantabrian Mountains, based on its disjunct distribution relative to typical European *Q. robur*, its occurrence in high-mountain habitats, and morphological features such as longer petioles and distinctive cupule scales (Rivas-Martínez *et al.*, 2002; Vila-Viçosa *et al.*, 2020a). In our sampling design, *Q. broteroana* and *Q. orocantabrica* were therefore initially treated as distinct field and literature-based sampling labels, and both names were retained in the analytical matrices, maps and figures for traceability. However, the present phylogenomic analyses do not support their recognition as independent evolutionary lineages. Samples assigned to *Q. orocantabrica* fell within the genomic variation of *Q. broteroana* in the phylogenetic and population-structure analyses. We therefore interpret both initial labels as representing a single Northwestern Iberian *Q. robur s.l.* lineage. The nomenclatural consequences of this result require a separate formal treatment; in the present work we retain both names only to document the correspondence between the original taxonomic assignments and the genomic entity recovered in this study.
If the Northwestern Iberian lineage recovered here is treated at species rank, the correct name should be evaluated through a formal nomenclatural assessment of the available names, basionyms, ranks and types, including *Q. robur* subsp. *broteroana* O.Schwarz and *Q. orocantabrica* Rivas Mart., Penas, T.E.Díaz & Llamas. We therefore avoid making a new nomenclatural act in the present phylogenomic study.

**Differential morphological traits:** In comparison with typical European *Q. robur* L., the Northwestern Iberian pedunculate-oak lineage (i.e. *Quercus broteroana* / *Quercus orocantabrica* complex) tends to have thicker and more coriaceous leaf blades, usually oblong to oblong-transovate, often broader and glossy on the adaxial surface. Leaves commonly show unequal lobes and more then 6–8 secondary veins, which are often sinuate. Cupules are relatively large, usually 18–25 mm, with brownish scales that are larger and freer near the base, becoming smaller and more appressed towards the apex. Petioles and peduncles also tend to be longer than in typical *Q. robur*. The Cantabrian material described as *Q. orocantabrica* shares this general morphology, especially the larger petioles and cupules, but has also been described as having pubescent, reddish cupule scales (Schwarz, 1937; Rivas-Martínez & Saénz, 1991; Rivas-Martínez *et al.*, 2002)

**Subsection *Roburoides*** (Schwarz, 1936a)

***Quercus pauciradiata*** Penas, Llamas, Pérez Morales & Acedo

**General information:** Taxon was another narrow endemic species described for the León Province, which could be segregated from surrounding roburoid oaks, namely *Q. petraea* and *Q. pyrenaica*, by having exclusive fasciculate trichomes and distinguishable leaf phenology and morphology (Penas *et al.*, 1997). The status of *Q. pauciradiata* may be interpreted in parallel with the Northwestern Iberian *Q. robur s.l.* lineage recovered in the present study. Just as the *Q. broteroana*–*Q. orocantabrica* complex appears to represent a differentiated Iberian expression of the broader pedunculate-oak assemblage, *Q. pauciradiata* may represent a southern Iberian expression of the *Q. petraea s.l.* complex. This interpretation is also relevant to the historical *Q. huguetiana* concept, although the latter requires dedicated typological, morphological and genomic reassessment before any nomenclatural conclusion can be drawn.

**Differential morphological traits:** Lobate to pinnatifid leaves, with different foliation period among sympatric species (*Q. petraea* and *Q. pyrenaica*), and fasciculate shorter trichomes with 2-4 rays forming a "cross-like" pattern (Penas *et al.*, 1997).

**Subsection *Galliferae*** Gürke (1897)

***Quercus canariensis*** Willd.

**General information:** Taxon is a relictual tree, with five subpopulations in the Iberian Peninsula. In spite of being largely distributed above limestones in North Africa, in the Iberian Peninsula it is related with silicious bedrock in places with mesophytic conditions, and has a higher demand on annual and summer precipitations in comparison with all *Galliferae* species (Vila-Viçosa *et al.*, 2015; Vila-Viçosa *et al.*, 2020a; Vila-Viçosa *et al.*, 2020b). It is known to form hybrid swarms with other sympatric oaks like *Quercus faginea* Lam. (*Q. ×marianica* C. Vicioso (=*Q. ×tlemcenensis* Trab.)) and *Q. pubescens* Willd. (*Q. ×cerrioides* Willk. & Costa) (Battandier & Trabut, 1905; Vicioso, 1950; Vazquez *et al.*, 2020b). *Q. canariensis* has been a hallmark for conservation in the Iberian Peninsula, being

Critically Endangered in Portugal (Carapeto *et al.*, 2020), considering that they form peculiar Miocene forests with special conservation interest (Ojeda *et al.*, 2000).

**Differential morphological traits:** Glabrescent leaves obovate or progressively obovate towards the apex, crenate to serrate, sometimes with sinuate-lobulated margin; with short obtuse lobes, acute in young leaves, having a single indumentum of fasciculate-trichomes (sometimes reddish in dryer stations) that can detach, forming a cotton like fuzzy indumentum, mainly in the midrib and secondary veins, that are rectilinear (almost parallel and in higher number (>11)). Absence of stellate trichomes (present in the remaining *Galliferae* species (*Quercus faginea* and *Q. lusitanica*) (Willdenow, 1805; Rivas-Martínez & Saénz, 1991).

***Quercus faginea*** Lam.

**General information:** congregates the concepts of *Q. broteroi* and *Q. faginea*, usually considered as two independent taxa (Franco, 1990; Rivas-Martínez & Saénz, 1991). Following recent reviews of the type material for this taxon, it was concluded that the name formerly ascribed to the Western and Southern Iberian oak as *Q. broteroi* is in fact the original concept of *Q. faginea* (Vazquez *et al.*, 2020b). The taxon with North and Northeast distribution that can be considered as a different species should be addressed to the concept of *Q. muricata* Palau, as suggested by Villar (1958). *Q. faginea* has a broad distribution in submediterranean and Atlantic areas of the Iberian Peninsula and through North Africa up to Tunisia, with preference for base-rich soils, being common in limestone areas with an increase of annual or summer precipitation (Vila-Viçosa *et al.*, 2014; Vila-Viçosa *et al.*, 2020a).

**Differential morphological traits:** Oblong to obovate and ovate leaves, with crenate to serrate margin, sometimes denticulate and mucronated; tomentose to pubescent bellow, with stellate and multiradiate trichomes (Lamarck, 1785; Schwarz, 1964).

***Quercus lusitanica*** Lam.

**General information:** This taxon is a geoxylic shrub oak that occurs in the extreme western areas of the Iberian Peninsula and has a sub-population in Northern Africa (Morocco), presenting a mostly Atlantic distribution and not tolerating winter cold.

**Differential morphological traits:** Shrubs with oblong to obovate and ovate leaves (occasionally lobulated), with flat and almost sessile cuneate leaves with partially entire margins in the proximal third (Lamarck, 1785; Sampaio, 1910; Rivas-Martínez & Saénz, 1991).

**Subsection *Dascia*** (Kotschy) Schwarz (1936a)

***Quercus pyrenaica*** Willd.

**General information:** has a wide distribution throughout the submediterranean and temperate belts of the Iberian Peninsula. Being an edaphic indifferent, it is more common above siliceous bedrock, supporting summer drought when compared to *Q. broteroana* (Vila-Viçosa et al., 2020a).

**Differential morphological traits:** Crenate to pinnatifid heavily tomentose leaves, with fasciculate trichomes showing >0.5 mm rays.

***Quercus pubescens*** Willd.

**General information:** The taxon has its far-west European limit in northeast Iberian Peninsula, in spite of being the most widely distributed oak species in Europe. It is a typical submediterranean species that can occur in temperate areas (Vila-Viçosa *et al.*, 2020a), with preference for base-rich soils. The taxon has a complex history of infraspecific taxa in Southern Europe (Pasta *et al.*, 2016). It forms known hybrid swarms either with *Q. faginea* (*Q. ×subpyrenaica* Villar), or with *Q. canariensis* (*Q. ×cerrioides* Willk. & Costa).

**Differential morphological traits:** Crenate to pinnatilobed leaves, with tomentose nerves and with fasciculate trichomes (<0.3 mm) (Willdenow, 1796; Schwarz, 1936c; Schwarz, 1964).

**Hybrids**

***Quercus ×cerrioides*** Willk. & Costa

**General information:** Taxon originally described for the littoral Catalonian mountains, it was associated with *Q. pubescens* albeit with a leaf shape closer to broad *Q. cerris* Lam. (Willkomm, 1859). For this reason, this plant has been ascribed to be a hybrid between *Q. canariensis* and *Q. ×subpyrenaica*, due to the dentate-to-serrate leaf shape that suggests a *Galliferae* species as putative parental (besides the obvious participation of *Q. canariensis*) (Rivas-Martínez et al., 1991; Vázquez et al., 2018). However, our study confirms the most probable hypothesis, which is that *Q. pubescens* is the parental species besides *Q. canariensis*. Schwarz (1936c) and Willkomm and Lange (1861) traditionally address the name *Q. ×cerrioides* Willk. & Costa to the hybrids of *Q. pubescens* and *Q. faginea* (*Quercus ×subpyrenaica* Villar).

**Differential morphological traits:** Crenate to serrate and lobulated margins, commonly with truncated lobes resembling *Q. pubescens*, sharing the reddish fuzzy indument of *Q. canariensis* (Willkomm, 1859; Schwarz, 1936c), and sharing the fasciculate trichomes of *Q. pubescens*.

***Q. ×coutinhoi*** Samp.

**General information:** This taxon was described as a hybrid between *Q. faginea* and *Q. robur*. This was based on several heterogeneous materials retained in the LISU herbarium, related with plants earlier described by Coutinho (1888) and mostly collected from Sintra, Caldas da Rainha and Coimbra regions in Portugal. This hybrid raises concerns on the putative parental species, with both *Q. broteroana* and *Q. estremadurensis* as potential taxa.

**Differential morphological traits:** Highly polymorphic taxon, sharing characters of both *Galliferae* and *Robur* Sections (Schwarz, 1936a), with leaves tendentially presenting acute lobes. Leaves are glabrescent with scattered stellate trichomes in the abaxial surface. Shorter petioles and larger peduncles when compared to *Q. faginea*.

***Quercus ×duriensis*** Franco & Vasconcellos (≡*Quercus ×coutinhoi* f. *duriensis* Franco & Vasc.)

**General information:** This hybrid was originally described by Vasconcellos and Franco (1954) taking into account the *Q. robur s.l.* concept and a putative variety of *Q. faginea* (*Quercus lusitanica* f. *salicifolia* Cout.) that refers to *Q. faginea* samples from the Douro Basin in Portugal. The type location from where this hybrid was originally described concurs with the distribution of *Q. broteroana*. Samples of hybrids with *Q. faginea* were collected in one of these locations, where *Q. broteroana* was also sampled and confirmed through our molecular analysis.

**Differential morphological traits:** Highly polymorphic, with smaller leaves with crenate to lobated margins, showing uneven rounded lobes with glabrescent cuneate leaves, sparse stellate trichomes, and short peduncles.

***Quercus faginea × Q. pyrenaica***

**General information:** This hybrid was traditionally named after *Q. ×neomairei* A. Camus (Franco, 1990), albeit studies of the Iberian white oaks' type material (Vila-Viçosa *et al.*, 2023) allowed to conclude that the specimen from the original type material kept in LISU of *Q. ×neomairei*, was instead a hybrid of *Q. pyrenaica* with *Q. estremadurensis* rather than a hybrid of *Q. pyrenaica* with *Q. faginea*. Taking this into account we used the unnamed designation for this taxon.

**Differential morphological traits:** Crenate to almost pinnatifid leaf margins, normally with deeper cut lobes. Heavily tomentose with a double indumentum of both fasciculate and stellate trichomes.

***Quercus ×fontqueri*** O.Schwarz

**General information:** This hybrid was described by Schwarz (1936c) as the natural hybrid between *Q. canariensis* and *Q. pyrenaica*, collected in Catalonia. Hybrids from southwest Portugal were identified in locations where parentals were present.

**Differential morphological traits:** Wide leaves with a pinnately lobed margin, and both 1) yellow-orangish fuzzy indumentum of *Q. canariensis* near the midrib and axilary joints with secondary nerves, and 2) fasciculate tricomes of *Q. pyrenaica* across the abaxial surface.

***Quercus ×marianica*** C.Vicioso (=*Q. ×tlemcenensis* Trab.)

**General information:** This hybrid is mostly distributed in southwestern Iberian Peninsula, in areas of mixed stands where *Q. canariensis* contacts with *Q. faginea*, normally in siliceous bedrocks (Vicioso, 1950; Vila-Viçosa *et al.*, 2015).

**Differential morphological traits:** Lanceolate leaves normally with rectilinear nerves (>11). Leaves either glabrescent with scarce stellate trichomes (*Quercus ×marianica* C.Vicioso), or tomentose with both stellate and fuzzy-like indumentum of fasciculate trichomes (*Q. ×tlemcenensis* Trab.) (Battandier & Trabut, 1905; Vicioso, 1950).

***Quercus ×subpyrenaica*** Villar

**General information:** It forms a continuous hybrid swarm in the Pre-Pyrenees mountains, with *Q. pubescens* and *Q. faginea* as parental species (Villar, 1935; Villar, 1958). Has a typical transitional submediterranean distribution (Himrane *et al.*, 2004; Vila-Viçosa *et al.*, 2020a).

**Differential morphological traits:** Lobate to dentate and serrate margins, with both stellate and fasciculate trichomes, the former notoriously visible in the midrib (Villar, 1935; Villar, 1958; Rivas-Martínez & Saénz, 1991).

**Note S2** Natural-history and historical-taxonomic context for Eurasian white-oak evolution

The interpretation of Eurasian white-oak evolution benefits from integrating phylogenomic evidence with historical taxonomy, natural history, and classical herbarium-based observations. Several classical infrageneric systems already recognised a deep heterogeneity within Eurasian white oaks, separating temperate roburoid complexes from southern, marcescent, pubescent, and eastern Mediterranean lineages later associated with Dascia, Macrantherae, Lanuginosae, and related concepts (Camus, 1935; Maleev, 1935; Schwarz, 1936a; Schwarz, 1936c; Schwarz, 1936b; Menitsky, 1972). Several patterns recovered in our molecular analyses are broadly consistent with earlier taxonomic treatments that organised white-oak diversity according to geography, indumentum, leaf persistence, marcescence, and hybridisation-prone contact zones, rather than by isolated diagnostic characters (Schwarz, 1935; Schwarz, 1936b; Schwarz, 1936a; Schwarz, 1936c; Vicioso, 1950; Villar, 1958). Although some of these historical infrageneric units are no longer followed as formal classifications, they provide useful biogeographic and morphological context for interpreting the structure, reticulation, and regional differentiation observed in the present study. These historical interpretations also provide a useful framework for reading the molecular signal below the sectional level, especially where classical taxa anticipated geographically-coherent lineages rather than merely typological species.

Several southern European taxa historically placed around *Q. robur s.l.* and *Q. petraea s.l.* show intermediate character combinations that contributed to long-standing ambiguity between roburoid, petraeoid, and locally differentiated southern European lineages (Schwarz, 1935; Schwarz, 1937; Camus, 1939; Vicioso, 1950; Menitsky, 1976, 1977). This is particularly relevant because several names historically treated within the broad *Q. robur–Q. petraea* complex may present intermediate morphological features, pending on geography. Specifically, names traditionally associated with *Q. robur s.l.*, including *Q. broteroana*, *Q. brutia*, *Q. estremadurensis*, *Q. haas*, *Q. hartwissiana*, *Q. imeretina*, and *Q. pedunculiflora*, were repeatedly segregated on combinations of cupule, peduncle, petiole, leaf texture, venation, and lobe architecture, but their rank and circumscription remained unstable across treatments (Schwarz, 1935; Schwarz, 1937; Camus, 1939; Vicioso, 1950; Menitsky, 1976, 1977). Meanwhile, several eastern or southern taxa associated with *Q. petraea s.l.*, including *Q. abietum*, *Q. cedrorum*, and *Q. mannifera*, show how the petraeoid complex also extends into Mediterranean and Near Eastern taxonomic problems that cannot be resolved from morphology alone (Camus, 1939; Menitsky, 1976, 1977). These historical concepts underline the need for broader molecular characterization across the biogeographical range of southern Europe, ideally based on bona fide taxonomic representatives collected from type populations, neighbouring areas, and contact zones.

The case of *Q. estremadurensis* is particularly relevant in this context. This taxon occupies a peripheral position along the western Iberian and submediterranean fringe, and has long been interpreted as a singular element within *Q. robur s.l.*. Its original treatment as an independent species, followed later reduction or absorption into *Q. robur s.l.* concepts, exemplifies the difficulty of assigning peripheral Iberian roburoid populations to either local relictual lineages or geographic variants of widespread European taxa (Schwarz, 1935; Schwarz, 1964; Camus, 1939; Vicioso, 1950). Its placement highlights the potential evolutionary relevance of southern European peninsulas as areas where temperate white-oak lineages may have persisted, differentiated, or retained relictual genomic variation. Broader sampling of southern European roburoid and petraeoid taxa will be necessary to clarify whether such taxa represent local derivatives of widespread lineages, remnants of older diversification processes, or geographically structured components of a wider European white-oak syngameon.

Historical taxonomy is especially informative for interpreting the Mediterranean and submediterranean white-oak complex, where classical systems repeatedly grouped taxa along west–east Mediterranean, Anatolian, Caucasian, and Near Eastern axes (Kotschy, 1862; Schwarz, 1936a; Camus, 1935; Maleev, 1935; Menitsky, 1976, 1977). The historical placement of *Q. pyrenaica*, *Q. frainetto*, *Q. macranthera*, *Q. vulcanica*, *Q. kotschyana*, *Q. canariensis*, *Q. boissieri*, and *Q. pubescens* in southern or eastern white-oak complexes anticipated a biogeographic signal that is partly recovered by the molecular evidence of the present work (Schwarz, 1936a; Schwarz, 1936c; Camus, 1935; Maleev, 1935; Menitsky, 1976, 1977). The taxa historically associated with the *Dascia–Macrantherae* problem are not recovered here as a simple typological group; rather, *Q. pyrenaica*, *Q. kotschyana*, *Q. vulcanica*, *Q. macranthera*, *Q. frainetto*, and related taxa are better interpreted as geographically structured components of a wider submediterranean and eastern Mediterranean assemblage (Kotschy, 1862; Schwarz, 1936a; Camus, 1935; Maleev, 1935; Menitsky, 1976, 1977). Near Eastern, Anatolian, and Caucasian species derive from a well-known hotspot of oak radiation and diversity (Petit *et al.*, 2002; Tarkhnishvili *et al.*, 2012; Schirone *et al.*, 2014). Camus also grouped several of these taxa in a common subsection, *Macrantherae*, relating *Q. macranthera* with *Q. frainetto* and *Q. pyrenaica* (Camus, 1935), while comparable eastern and Caucasian affinities were also emphasised in later Eurasian treatments of white oaks (Maleev, 1935; Menitsky, 1972). In this context, the relationships among *Q. macranthera*, *Q. vulcanica*, *Q. kotschyana*, *Q. boissieri*, and *Q. pyrenaica*, signalled in the present work as Group B species, fit the evolutionary expression of a geographically structured submediterranean assemblage, shaped by either shared ancestry, regional isolation, morphological convergence, or secondary contact.

The biogeographic relationships of the *Q. canariensis–Q. boissieri* complex are also relevant to the interpretation of Mediterranean white-oak differentiation. The proximity between *Q. canariensis* and *Q. boissieri* gives renewed relevance to their historical placement in Series Orientales, which was supported by shared foliar architecture and by the presence, in both taxa, of dense fasciculate trichomes forming a cottony indumentum along the abaxial veins (Schwarz, 1936a, 1936b). Rather than supporting a simple east–west vicariant pair, our results support a broader Mediterranean structuring scenario, in which *Q. canariensis* and *Q. boissieri* occupy opposite margins of the complex, while *Q. pubescens* represents a geographically intermediate and reticulate component of Group B species(Humphries *et al.*, 2017).

The position of *Q. pubescens* is particularly relevant because it was historically difficult to accommodate within a strict *Q. robur–Q. petraea* framework. Its placement among pubescent and southern marcescent oaks, rather than within a narrow roburoid concept, is consistent with earlier treatments that linked *Q. pubescens* to southern European and eastern Mediterranean white-oak complexes (Schwarz, 1936c; Camus, 1935; Menitsky, 1972). Its broad submediterranean distribution and sympatry with several related oaks, including *Q. faginea, Q. canariensis, Q. boissieri*, and *Q. petraea*, make it a key taxon for interpreting the transition between temperate roburoid oaks and the southern submediterranean–*Macrantherae* (Group B) assemblage recovered in the present study.

Within the western Mediterranean, the submediterranean ecotone is especially important for understanding the differentiation of Iberian and neighbouring white oaks. This interpretation is consistent with treatments that emphasised geography, ecological continuity, and transitional morphologies in the western Mediterranean *Q. faginea s.l.* complex, rather than excessive splitting into narrowly defined groups (Villar, 1935; Villar, 1958; Vicioso, 1950). This transitional ecotone may have contributed to the evolutionary history of Mediterranean Group B oaks by linking thermophilic conditions with lower precipitation seasonality, partially resembling subtropical conditions inferred for the late Miocene. Such conditions are commonly associated with areas where relictual lauroid and broad-leaved flora persisted (Suc, 1984; Rodríguez-Sánchez *et al.*, 2010; Rodríguez-Sánchez & Arroyo, 2011; Vila-Viçosa *et al.*, 2020c). In this sense, the Iberian Peninsula should not be interpreted only as a terminal refuge, but also as a structured contact region where ecological gradients, historical persistence, and hybridization have interacted through time.

The hybrid complexes documented in the Iberian Peninsula further support the biological relevance of classical nothotaxonomic treatments. Several nothotaxa, including Q*. ×cerrioides, Q. ×subpyrenaica, Q. ×marianica, Q. ×fontqueri, Q. ×coutinhoi, Q. ×duriensis*, and *Q. faginea × Q. pyrenaica*, correspond to historically described or morphologically recognized entities that are consistent with reticulate patterns recovered in the present study, especially in the western and northeastern Iberian contact zones where *Q. faginea*, *Q. canariensis*, *Q. pyrenaica*, *Q. pubescens*, and roburoid taxa overlap (Coutinho, 1888; Schwarz, 1936c; Vicioso, 1950; Villar, 1958; Rivas-Martínez & Sáenz, 1991). In particular, the northeastern Iberian hybrid swarm involving *Q. pubescens* and *Q. canariensis* and signalled by our molecular study, clarifies the interpretation of *Q. ×cerrioides*, historically described as stemming from the Catalonian littoral mountains, and subsequently interpreted through different parental combinations involving *Q. pubescens*, *Q. faginea*,

*Q. ×subpyrenaica*, and *Q. canariensis* (Willkomm, 1859; Willkomm & Lange, 1861; Schwarz, 1936c; Rivas-Martínez & Sáenz, 1991; Rivas-Martínez et al., 2002). This supports the interpretation of northeastern Iberia as a reticulate contact zone between western Mediterranean and central European submediterranean white-oak lineages, rather than as a simple extension of the *Q. faginea* hybrid complex.

Overall, the convergence between historical taxonomy and molecular evidence reinforces the need to interpret Eurasian white-oak evolution as the result of simultaneous macroevolutionary and microevolutionary processes. Broad geographic structuring, old biogeographic disjunctions, ecological differentiation, local persistence, and recurrent hybridization all contribute to the genomic patterns observed in Iberian and Mediterranean white oaks. In this sense, the Iberian Peninsula is not only a refuge or a peripheral reservoir of white-oak diversity, but a phylogenomically structured contact region where roburoid, submediterranean, and Macrantherae-related lineages meet, differentiate, and hybridise. Future studies should combine dense phylogenomic sampling, type-locality material, herbarium revision, and explicit sampling of contact zones and known hybrid swarms, including those retrieved from grey historical literature. Such integration is essential for distinguishing taxonomic artefacts from evolutionarily meaningful lineages and for understanding the role of southern European peninsulas in the diversification of Western Palearctic white oaks.

Kotschy T. 1862. *Die Eichen Europa's und des Orient's*. Wien und Olmütz: E. Hölzel.

Lamarck JB. 1785. *Encyclopédie méthodique. Botanique*. Paris: Panckoucke.

Loidi J. 2017. *The vegetation of the Iberian Peninsula*: Springer.

Maleev VP. 1935. A Review of Oak Trees in Caucasus through theirs Taxonomical and geographical relations and concerning to evolution of group Robur. *Bot. Zhur.* 20(2): 156-172.

Médail F, Baumel A. 2018. Using phylogeography to define conservation priorities: The case of narrow endemic plants in the Mediterranean Basin hotspot. *Biological Conservation* 224: 258-266.

Menitsky G. 1972. *Generis Quercus L. species asiae austro-occidentalis*.

Ojeda F, Marañón T, Arroyo J. 2000. Plant diversity patterns in the Aljibe Mountains (S. Spain): a comprehensive account. *Biodiversity & Conservation* 9(9): 1323-1343.

Pasta S, de Rigo D, Caudullo G 2016. Quercus pubescens in Europe: distribution, habitat, usage and threats. In: San-Miguel-Ayanz J, de Rigo D, Caudullo G, Houston Durrant T, Mauri A eds.

*European atlas of forest tree species*. Luxembourg: Publications Office of the European Union, 156-157.

Penas A, Llamas F, Perez-Morales C, Acedo C. 1997. A new species of Quercus L. (Fagaceae) from NW Spain. *Botanica Helvetica* 107: 75-82.

Petit RJ, Brewer S, Bordács S, Burg K, Cheddadi R, Coart E, Cottrell J, Csaikl UM, van Dam B, Deans JD. 2002. Identification of refugia and post-glacial colonisation routes of European white oaks based on chloroplast DNA and fossil pollen evidence. *Forest ecology and management* 156(1): 49-74.

Rivas-Martínez S, Báscones J.E., Díaz T., Fernández-González F., J. L. 1991. Vegetación del Pirineo occidental y Navarra. *Itinera Geobotanica* 5: 530.

Rivas-Martínez S, González-Días TE, González FF, Loidi JJ, Lousã MF, Merino ÁP. 2002. Vascular plant communities of Spain and Portugal: addenda to the syntaxonomical checklist of 2001. Part II. *Itinera Geobotanica*(15): 433-922.

Rivas-Martínez S, Penas A, del Río S, González T, Rivas-Sáenz S 2017. Bioclimatology of the Iberian Peninsula and the Balearic Islands. In: Loidi J ed. *The Vegetation of the Iberian Peninsula*. Utrecht: Springer, 29-80.

Rivas-Martínez S, Saénz C. 1991. Enumeración de los Quercus de la Península Ibérica. *Rivasgodaya* 6: 101-110.

Rodríguez-Sánchez F, Arroyo J. 2011. Cenozoic climate changes and the demise of Tethyan laurel forests: lessons for the future from an integrative reconstruction of the past. *Climate change, ecology and systematics*: 280-303.

Rodríguez-Sánchez F, Hampe A, Jordano P, Arroyo J. 2010. Past tree range dynamics in the Iberian Peninsula inferred through phylogeography and palaeodistribution modelling: a review. *Review of Palaeobotany and Palynology* 162(3): 507-521.

Sampaio G. 1910. Nota a propósito dos Quercus lusitanica Lamk., Q. humilis Lamk e Quercus faginea, Lamk. . *Annaes Scientificos da Academia Polytechnica do Porto* 5(3): 161-165.

Schirone B, Spada F, Piovesan G, Simeone MC 2014. Phenorhythms and forest refugia. *Warm-temperate deciduous forests around the Northern hemisphere*: Springer, 213-223.

Schwarz O. 1935. Einige neue Eichen des Mediterrangebiets und Vorderasiens. *Notizblatt des Königl. botanischen Gartens und Museums zu Berlin* 114: 461-469.

Schwarz O. 1936a. Entwurf zu einem natürlichen System der Cupuliferen und der Gattung Quercus L. *Notizblatt des Botanischen Gartens und Museums zu Berlin-Dahlem*: 1-22.

Schwarz O. 1936b. *Monographie der Eichen Europas und des Mittelmeergebietes*: Selbstverlag.

Schwarz O. 1936c. Sobre los Quercus catalanes del subgénero Lepidobalanus Oerst. *Cavanillesia* 8: 65–100.

Schwarz O. 1937. Monographie der Eichen Europas und des Mittelmeergebietes. *Repertorium novarum speciarum regni vegetabilis* 1: 1-200.

Schwarz O 1964. Quercus L. In: Tutin TG, Heywood VH, Burges NA, Valentine DH, Walters SM, DA. W eds. *Flora Europaea*. Cambridge: Cambridge University Press, 61-64.

Schwarz O 1993. Quercus L. In: al. TTe ed. *Flora Europaea*. Cambridge: Cambridge University Press., 72-76.

Suc J-P. 1984. Origin and evolution of the Mediterranean vegetation and climate in Europe. *Nature* 307(5950): 429-432.

Tarkhnishvili D, Gavashelishvili A, Mumladze L. 2012. Palaeoclimatic models help to understand current distribution of Caucasian forest species. *Biological journal of the Linnean Society* 105(1): 231-248.

Vasconcellos JC, Franco JA. 1954. Os Carvalhos de Portugal. *Anais do Instituto Superior de Agronomia* 21: 1–135.

Vazquez FM, Coombes A, Almeida R, Garcia-Alonso D, Marquez-Garcia F, Vila-Vicosa C. 2020a. Lectotypification of names of Quercus spp.(Fagaceae) described by Lamarck from the Iberian Peninsula. *Phytotaxa* 455(3): 205–213-205–213.

Vazquez FM, Coombes A, Almeida R, Garcia-Alonso D, Marquez-Garcia F, Vila-Vicosa C. 2020b. Lectotypification of names of Quercus spp.(Fagaceae) described by Lamarck from the Iberian Peninsula. *Phytotaxa* 455(3): 205-213.

Vázquez FM, Coombes AJ, García D, Márquez F, Meireles C, Barrena MJ, Vila-Viçosa C. 2018. Anotaciones a la nomenclatura del género Quercus L., (FAGACEAE) en la Península Ibérica y NW de África. . *Folia Botánica Extremadurensis* 12: 5-79.

Vicioso C. 1950. *Revisión del género Quercus en España*. Madrid: Tipografía Artítica.

Vila-Viçosa C, Arenas-Castro S, Marcos B, Honrado J, García C, Vázquez FM, Almeida R, Gonçalves J. 2020a. Combining Satellite Remote Sensing and Climate Data in Species Distribution Models to Improve the Conservation of Iberian White Oaks (Quercus L.). *ISPRS International Journal of Geo-Information* 9(12): 735.

Vila-Viçosa C, Capelo J, Alves P, Almeida R, Vázquez FM. 2023. New annotated checklist of the Portuguese oaks. *Mediterranean Botany* 44: 1-32.

Vila-Viçosa C, Gonçalves J, Honrado J, Lomba Â, Almeida RS, Vázquez FM, Garcia C. 2020b. Late Quaternary range shifts of marcescent oaks unveil the dynamics of a major biogeographic transition in southern Europe. *Scientific reports* 10(1): 21598.

Vila-Viçosa C, Gonçalves J, Honrado J, Lomba Â, Almeida RS, Vázquez FM, Garcia C. 2020c. Late Quaternary range shifts of marcescent oaks unveil the dynamics of a major biogeographic transition in southern Europe. *Scientific reports* 10(1): 1-12.

Vila-Viçosa C, Vázquez F.M., Meireles C, Pinto-Gomes C. 2014. Taxonomic peculiarities of marcescent oaks (Quercus, Fagaceae) in southern Portugal. *Lazaroa* 35: 139–153.

Vila-Viçosa C, Vázquez FM, Mendes P, Del Rio S, Musarella C, Cano-Ortiz A, Meireles C. 2015. Syntaxonomic update on the relict groves of Mirbeck's oak (Quercus canariensis Willd. and Q. marianica C. Vicioso) in southern Iberia. *Plant Biosystems* 149(3): 512-526.

Villar EH. 1935. Sur le nom de quelques Quercus et la systématique du faginea. *Cavanillesia* 7: 57-70.

Villar EH. 1958. Estudios sobre los Quercus del Oeste mediterráneo. *Anales del Instituto Botanico de A.J. Cavanilles* 15: 3-114.

Willdenow CL. 1796. *Berlinische Baumzucht oder Beschreibung der in den Gärten um Berlin im Freien ausdauernden Bäume und Sträucher für Gartenliebhaber und Freunde der Botanik*. Berlin: Nauck.

Willdenow CL. 1805. *Species plantarum exhibentes plantas rite cognitas ad genera relatas cum differentiis specificis, nominibus trivialibus, synonymis selectis, locis naturalibus secundum systema sexuale digestas*. Berlin: Nauck.

Willkomm HM. 1859. *Pugillus plantarum novarum peninsulae pyrenaicae*: Journal für die Botanik.

Willkomm M, Lange J. 1861. *Prodromus florae Hispanicae. I.*: Stuttgart.
